# Prelimbic-dependent activation of amygdala somatostatin interneurons signals non-aversive cues to promote discrimination

**DOI:** 10.1101/2020.06.23.156018

**Authors:** J.M. Stujenske, P.K. O’Neill, I Nahmoud, S. Goldberg, L. Diaz, M. Labkovich, W. Hardin, S.S. Bolkan, T.R. Reardon, T.J. Spellman, C.D. Salzman, J.A. Gordon, E. Likhtik

**Affiliations:** Department of Psychiatry, Weill Cornell Medical College; Department of Neuroscience, Columbia University; Department of Chemistry, Hunter College, CUNY; Department of Biological Sciences, Barnard College, Columbia University; Department of Ecology, Evolution, and Environmental Biology, Columbia University; Department of Biological Sciences, Hunter College, CUNY; Department of Psychiatry, Columbia University; Princeton Neuroscience Institute, Princeton University; Brain and Mind Research Institute, Weill Cornell Medical College; National Institute of Mental Health; Department of Biological Sciences, Hunter College, CUNY; Biology Program, The Graduate Center, CUNY

## Abstract

The amygdala and prelimbic cortex (PL) communicate during fear discrimination retrieval, but how they coordinate to discriminate a non-threatening stimulus is unknown. Here, we show that somatostatin interneurons (SOM) in the basolateral amygdala (BLA) become active specifically during learned non-threatening cues, when they block sensory-evoked phase resetting of theta-oscillations. Further, we show that SOM activation is PL-dependent, and promotes discrimination of non-threat. Thus, fear discrimination engages PL-dependent coordination of BLA SOM responses to non-threatening stimuli.

---

Generalized fear is a cardinal symptom in disorders of trauma and anxiety [1], and is associated with a hyperactive amygdala and a hypoactive medial prefrontal cortex [2] [3] [4]. Neural signatures of fear discrimination are present in the basolateral amygdala (BLA) [5] [6, 7], and mice with dysfunctional BLA inhibitory circuits generalize fear [8] [9]. Parvalbumin-positive (PV) and somatostatin-positive (SOM) interneurons (INs) play opposite roles in the acquisition of fear conditioned stimuli [10] promoting and impeding fear acquisition, respectively. However, it is not known whether there is a specific role for BLA INs during discrimination of non-threat.

Discrimination of non-threat has yet to be conclusively demonstrated as an actively maintained process. Recent work showed that during conditioning, BLA principal neurons (PN) remap to encode fear conditioned stimuli, whereas encoding of neutral stimuli remains passively maintained [7]. However, another study found that discrimination learning recruits specific PNs and INs that prevent generalization [5]. Here, we sought to investigate how BLA INs participate during retrieval of learned non-threatening cues, and whether they are coordinated by upstream prefrontal activity, which is known to be a crucial region for fear discrimination learning.

We trained mice on a 3-day differential fear conditioning (DFC) paradigm [11] (Figure 1a). Mice were exposed to two tones (2kHz and 8kHz, 50ms pips presented at 1Hz for 30sec). One tone was randomly assigned to be paired with a shock (CS+), and the other was explicitly unpaired (CS-). After conditioning, mice were exposed to only the CS+ or the CS-, and BLA SOM and PV IN activity was quantified using c-Fos co-labeling (Figures 1b-d). From these two cell types, the SOM INs were significantly more active during the CS- (Figure 1c-d), although overall c-Fos expression (mostly capturing PNs) was not altered by CS type (Supplementary Figure 1), consistent with our prior electrophysiology findings [11, 12].

**Figure 1.**
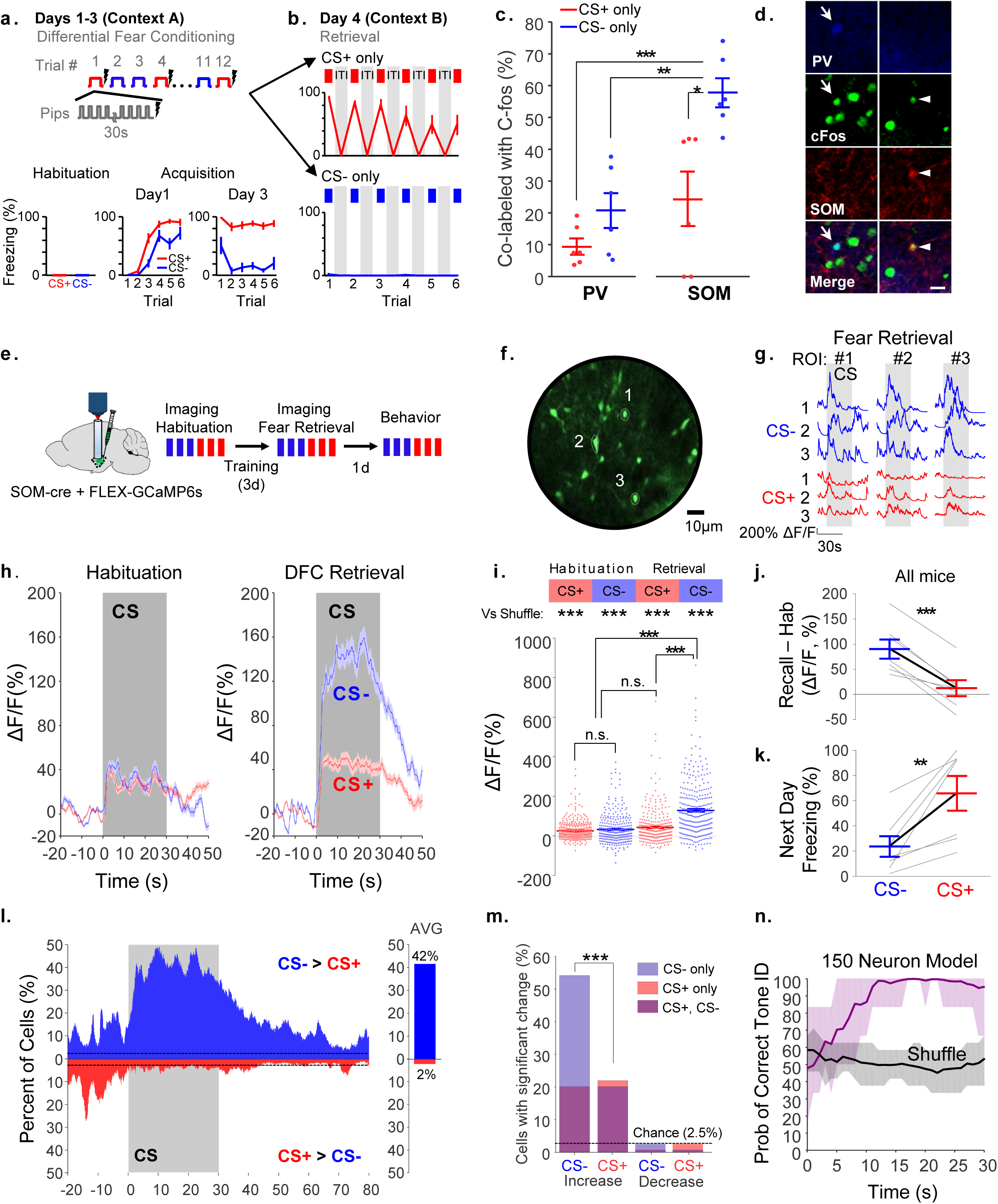
BLA SOM+ interneurons are selectively upregulated during CS-retrieval. **a.** Top, the differential fear conditioning paradigm includes 3 days of training, each with six presentations of the CS+ paired with shock, interspersed with six presentations of the CS-, explicitly unpaired with shock. CSs were 2kHz and 8kHz tones, counterbalanced between animals, that consisted of 50ms pips presented once per second for thirty seconds. 1 second foot shock began at the start of final pip. Bottom, mice (n=12) did not freeze to tones prior to fear conditioning. On the first day of DFC, mice freeze to both the CS+ and CS-, but by the third day of DFC, discrimination between the tones is already apparent, even though mice are still receiving foot shocks during the CS+. **b.** On day 4, mice were randomized to undergo CS+ (n=6) or CS- (n=6) retrieval, in the absence of shocks. On retrieval day, Mice froze significantly more to the CS+ (top, red) than to the CS- (bottom, blue), although freezing to the CS+ dropped over the six CS+ presentations, rm-ANOVA, CS: F_1,10_ =7.23, p<.0001, tone: F_5,50_=6.44, p=0.0001; CS x tone: F_5,50_, p=0.003. **c.** Mice were perfused 1.5 hours after fear retrieval, and the BLA was labeled with antibodies for parvalbumin, somatostatin, and cFos. The percentage of PV+ cells that were also cFos-co-expressing and the percentage of SOM cells that were also cFos-co-expressing was quantified for mice exposed to the CS+ (red) or CS- (blue). SOM INs show increased activity after CS-retrieval relative to all other conditions. Asterisks denote significant Bonferroni post-hoc test results. The unmarked pairs were not significant. One-way ANOVA, controlled for the effects of individual animals; F_2,10_=13.24, p=.0015. Post-hoc Bonferroni tests: CS+ PV vs CS- PV, p=0.58; CS+ PV vs CS+ SOM, p=0.51; CS+ PV vs CS- SOM, p=2.2 ×10^−5^; CS- PV vs CS- SOM, p=0.0045; CS- PV vs CS+ SOM, p=1.0; CS+ SOM vs CS- SOM, p=0.0042. **d.** Representative immunohistochemistry demonstrating a PV+ cell co-labeled with cFos (full arrow, left) and a SOM+ cell co-labeled with cFos (arrowhead, right). Scale bar, 20 μm. **e.** SOM-cre mice were injected with FLEX-GCaMP6s virus and SOM+ INs were imaged during habituation (3 CS-, 3 CS+), and fear retrieval (3 CS-, 3 CS+). Mice were exposed to another 3 CS- and 3 CS+ trials in a subsequent experiment (results shown in Figure 4), and then mice were tested for behavioral responses to CS- and CS+ the next day. **f.** Maximum projection image of example animal expressing GCaMP6s in BLA SOM+ INs. Three representative ROIs are shown in white contours. **g.** The CS- and CS+-evoked responses in the ROIs labeled in **f** are plotted for all CS presentations, demonstrating that SOM+ cells reliably responded to the CS- and weakly and irregularly responded to the CS+. Note that time advances as you progress down the figure. **h.** The average response of all SOM+ cells to the CS+ and CS-during habituation (left) and retrieval (right) are plotted as ΔF/F, normalized to the pre-tone period. **i.** The average response of each SOM+ cell to the CS+ and CS-. Asterisks next to “vs shuffle” refer to the statistical significance of the difference between cell responses to tones compared to that expected by chance (determined using a shuffled distribution, see methods). All tones evoked a significant response across the population. Asterisks showing between condition comparisons denote significant post-hoc Bonferroni corrected results. n.s., non-significant post-hoc results. CS- retrieval evokes higher SOM responses than other conditions. n=378 cells from 7 animals (habituation, n=322 cells (7 mice), retrieval, n=306 cells (7 mice), n=250 cells from both sessions). rmANOVA, Tone: F_1,875_=1.24, p=0.27; Session: F_1,875_=1.82, p=0.09; Tone x Session interaction, F_1,875_=76.16, p<0.001. Bonferroni post-hoc tests: CS- Hab vs CS+ Hab, p=0.6; CS- Hab vs CS- Retrieval, p = 5.3 × 10^−17^; CS- Hab vs CS+ Retrieval, p = 1.0; CS+ Hab vs CS+ Retrieval, p=0.08; CS+ Hab vs CS- Retrieval, p=7.1×10^−19^; CS- Retrieval vs CS+ Retrieval, p=3.7 × 10^−31^. **j.** The average change in response of all SOM+ cells within each individual animal from habituation to recall for CS- (blue) and CS+ (red) is plotted, with all mice demonstrating the same differential response, with a significantly stronger CS- response than CS+ response. **k.** The behavioral response of mice to the CS+ and CS- was quantified on day 6, after the mice had already been exposed to 6 CS+ and CS- tones during retrieval. Nevertheless, all mice froze more to the CS+ than CS-, confirming that they had appropriately acquired the differential fear conditioning learning, though at least some of the mice had evidence of extinguished response. **l.** (Left) The number of SOM+ cells with significantly greater response to the CS- than the CS+ is plotted on top in blue, and the number of cells with significantly greater response to the CS+ than CS- is plotted below in red (see methods). (Right) During the overall 30s tone, 42% of cells exhibited a statistically significantly greater response to the CS- than CS+, while only 2% of cells exhibited a significantly greater response to the CS+ (chance level: 2.5%). **m.** The percentage of SOM+ cells with significant response to the CS- (blue) and CS+ (red) is plotted for those that had an increase (left) or decrease (right) in their calcium response. Cells responding as increasers or decreasers to both the CS+ and CS- are plotted in purple, while the cells with selective response to only the CS+ and CS- are denoted in light purple or light red. Chance levels were set at 2.5% (dotted line). *** p < 0.001 McNemar’s test. **n.** Numerous 150 neuron models, subsampled from the full dataset across animals, were trained to decode CS+ and CS-. The probability of the model correctly determining tone identity at each second after tone onset is plotted as the mean model performance +/- 95% confidence intervals (purple), compared against results from models trained on data in which tone identity was shuffled (gray). Model performance steadily improved with time until plateauing, with significant decoding above chance levels by 10s after tone onset. *p<0.05, **p<0.01, ***p<0.001. Unless specified, data plotted as mean +/- SEM.

To further investigate SOM activation during the cues, we recorded the activity of BLA SOM INs using 2-photon imaging through a GRIN lens in SOM-cre mice injected with FLEX-GCaMP6s (Figure 1e-f). Responses of SOM INs to the CS+ and CS- were first imaged during stimulus habituation, and then again during retrieval after DFC (Figure 1g, Supplementary Figure 2). SOM INs had a weak response to both stimuli during habituation (Figure 1h). After DFC, there was a large increase in CS-response without a change in CS+ response (Figure 1h-j), and all mice demonstrated discrimination during freely-moving recall on the next day (Figure 1k). Notably, we only used calcium imaging data collected during periods of immobility for analysis such that results did not merely reflect locomotion. Individual SOM INs responded selectively to the CS-, with over 40% of cells showing a significantly stronger response to CS- than CS+ (Figure 1l, see methods), whereas only 2% responded more to the CS+. Overall, 54% of SOM INs were significantly active to the CS-compared to 22% to the CS+ (of which 91% were also responsive to the CS-, Figure 1m). The preferential CS-response was evident within 5 seconds of tone onset (Figure 1l, Supplementary Figure 3). Principle component analysis showed that the CS-response was relatively homogeneous (81% of variance explained by Principle Component 1, which correlated with the average fluorescence response, R=0.98, p<.001). We confirmed that this differential response was sufficient to encode tone identity using a linear classifier (support vector machine, see methods). Decoding success steadily improved over the first 10s of tone presentation, plateauing at 95-100% accuracy, confirming a significant differential response to CS-, compared to CS+, in the SOM population in all animals (Figure 1n, Supplementary Figure 4).

Next, we investigated whether fear discrimination specifically depends on BLA SOM Ins or if PV INs also affect discrimination. SOM or PV cells were optogenetically silenced during DFC retrieval by delivering light bilaterally to the BLA on half of the tone presentations in SOM-cre and PV-cre mice expressing cre-dependent eArch or control virus in the BLA (Figure 2a-d, Supplementary Figure 5). Electrodes were also implanted for concurrent electrophysiological recordings.

**Figure 2.**
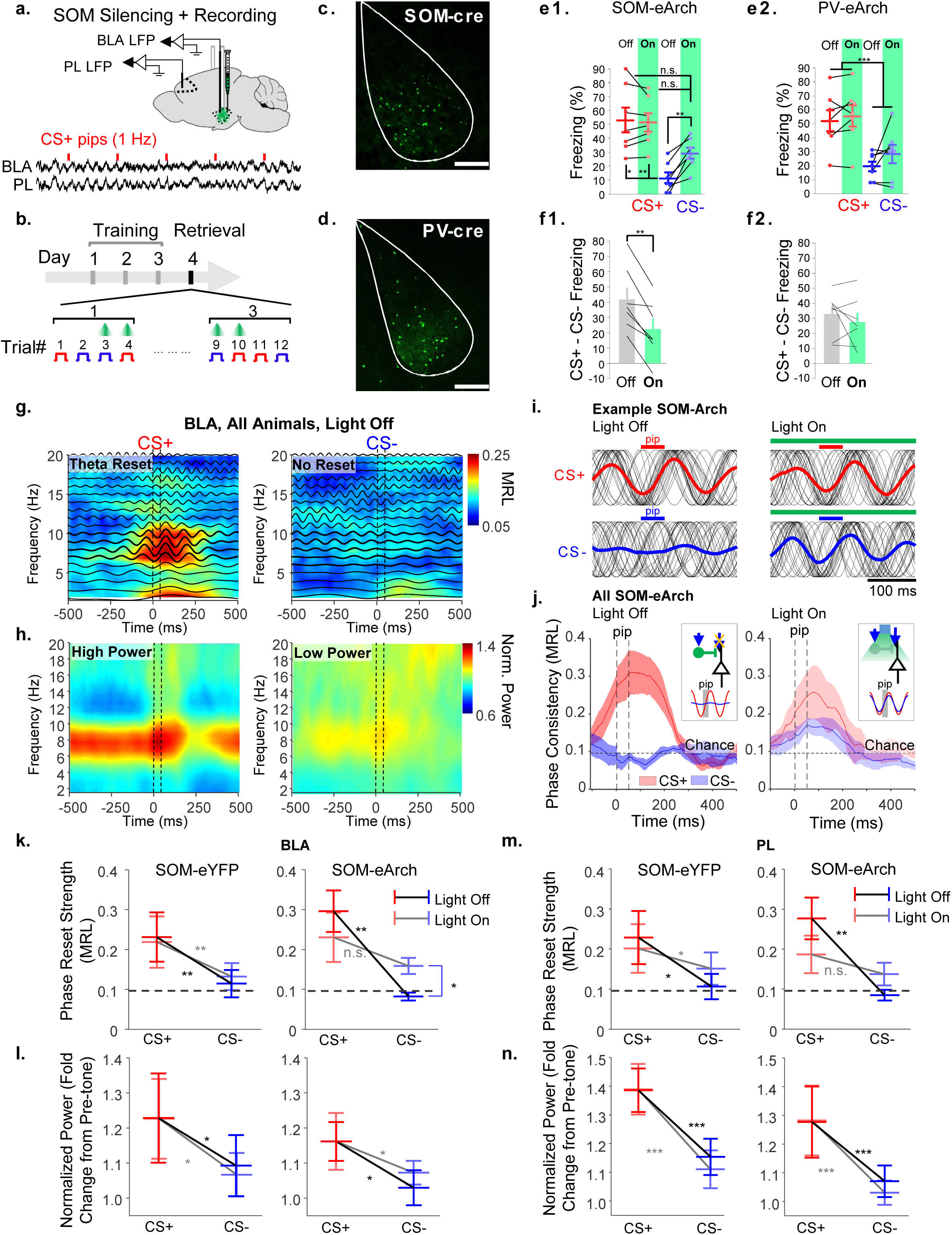
BLA SOM+ interneurons mediate discrimination by desynchronizing inputs to the amygdala during non-threatening stimuli. **a.** Cre-dependent AAV2/5 viruses expressing either eArchaerhodospin3.0-eYFP or eYFP were injected into the BLA of SOM-cre and PV-cre animals. A subset of SOM-cre mice expressing FLEX-eYFP or FLEX-eArch3.0 virus also had electrodes implanted in the BLA and PL for recording local field potentials simultaneously with optogenetic manipulation. Top, schematic of experiment. Bottom, example LFP recordings from the BLA and PL of a representative animal during one CS+ presentation. Red lines indicate the times of CS+ pip presentations. **b.** The mice were trained for three days on DFC and on the fourth day, they were exposed to 3 CS+s paired with light, 3CS+s without light, 3CS-s with light, and 3CS-s without light, which were interleaved pseudorandomly in a block-wise fashion (3 blocks containing each of the four conditions). **c and d.** Representative histology of SOM-cre (c) and PV-cre (d) animals expressing FLEX-eYFP virus in the BLA. Note an absence of central amygdala expression. Scale bars, 200 µm. **e.** Percent freezing of SOM-cre (**e1**) and PV-cre (**e2**) animals expressing FLEX-eArch3.0 in response to CS+ (red) and CS- (blue) in the presence (lighter colors) and absence (darker colors) of fiber optic illumination of the BLA. In **e1**, asterisks denote Bonferroni post-hoc test results, whereas in **e2**, asterisks denote CS effect, with insignificant effect of light or CS x light interaction. **e1.** SOM-cre + FLEX-eArch, n=7, rm-ANOVA: CS: F_1,18_=49.08, p<0.001; Light: F_1,18_=3.18, p=.09; CS x light interaction: F_1,18_=4.52, p=0.048; post hoc Bonferroni tests: CS+ Off vs CS+ On, p=1.0; CS+ Off vs CS- Off, p=0.011; CS+ Off vs CS- On, p=0.20; CS- Off vs CS- On, p=0.0053; CS+ On vs CS- On, p=0.12; CS- Off vs CS+ on; p=0.0052. **e2.** PV-cre + FLEX-eArch, n=7, rm-ANOVA: CS: F_1,18_=57.08, p<0.0001; light: F_1,18_=2.43, p=0.1364; CS x light interaction: F_1,18_=0.44, p=.52. Bonferroni-corrected paired t-test for Difference in Freezing, Light Off vs On: p=0.69. **f.** Difference between CS+ and CS- freezing during light off (gray) and light on (green) for SOM-cre (**f1**) or PV-cre (**f2**) mice, as in **e**. Bonferroni-corrected (for 4 comparisons including comparisons for eYFP-expressing animals, found in Supplementary Figure 5) paired t-tests, Light Off vs On: SOM-eArch, p=0.0099; PV-eArch, p=0.69. *p<0.05, **p<0.01, ***p<0.001. All data plotted as mean +/- SEM. **g.** The phase consistency of oscillations recorded in the BLA LFP, averaged over all SOM-cre animals (n=13; n=6 expressing FLEX-eArch3.0 and n=7 expressing FLEX-eYFP) during CS+ (left) and CS- (right), in the absence of fiber optic light delivery. Phase consistency is plotted as a heat map for different frequency bands at different times relative to pip onset, with warmer colors indicating stronger phase reset. Phase consistency is quantified as the mean resultant length (methods and Supplementary Figure 6). The average pip-evoked signal in different frequency bands (2 Hz bandwidth, centered at frequency on y axis) are overlaid on top of the heat map. A phase reset is seen in the 6-10 Hz frequency band during CS+ presentations but not CS- presentations. **h.** The average power across animals (as in **g**, n=13) during CS+ (left) and CS- (right), in the absence of fiber optic light delivery, is plotted as a spectrogram. Power is elevated in the 6-10 Hz theta frequency band during the CS+ but not the CS-. **i.** All pip-evoked theta (6-10 Hz) filtered LFPs (black traces) are overlayed from a representative Arch-expressing SOM-cre mouse during CS+ (red) and CS- (blue) in the presence and absence of SOM+ IN silencing (Light Off and Light On). These signals have been made “phase pure,” equalizing the amplitude, for illustrative purposes. Averages are plotted as overlying red and blue traces. The pips are plotted as red and blue rectangles. **j.** Average theta (6-10 Hz) phase consistency of all Arch-expressing animals (n=6) over time is plotted for the CS+ (red) and CS- (blue) during Light Off (left) and Light On (right) trials. Silencing of SOM+ INs induces a theta reset during the CS-, as shown by an elevation of phase consistency above chance, similar to the CS+ (statistics in **k**). *Inset (Left):* A schematic illustrating the effect: when the CS- (blue arrow) is presented, SOM INs (green cell) are activated and block theta reseting via dendritic inhibition on principal cells. This allows for lower input-driven synchronous activation of BLA PNs (white cell), and lower theta reset (blue line) compared to the CS+ (red line), when SOM INs are quiet, allowing for synchronous PN activation by incoming input (grey box denotes auditory pip). *Inset (Right)*: When SOM INs are silenced, sensory inputs drives PNs during the CS- (blue arrow), and induces a higher theta reset (blue line), similar to the CS+ in the presence of a pip. Note that the signal shown in the *Insets* is a sine wave to scale the magnitude of the average resets, not a representative example, which is shown in **i**. **k.** Average pip-evoked theta (6-10 Hz) phase consistency in the BLA during the CS+ (red) and CS- (blue) in SOM-cre mice expressing eYFP (left) or eArch (right) in the presence or absence of light. As shown in **j**, silencing SOM+ INs induces a theta reset to the CS-. The artificial CS-induced theta reset was still smaller in magnitude than during the CS+, though this difference did not reach statistical significance (see below). Astericks for eYFP indicate the main effect of CS, while notations for eArch reflect significant Bonferroni post-hoc test results. **eYFP**, n=7, rm-ANOVA: CS: F_1,18_=13.24, p=0.0020; Light: F_1,18_=0.02, p=0.90; CS x Light interaction: F_1,18_=0.9, p=0.36. **eArch**, n=6, rm-ANOVA: CS: F_1,15_=15.41, p=0.0014; Light: F_1,15_=0.48, p=0.50; CS x Light interaction: F_1,15_=8.9, p=0.0093. Bonferroni post-hoc tests: CS+ Off vs CS+ On, p=0.42; CS+ Off vs CS- Off, p=0.0043; CS+ Off vs CS- On, p=0.36; CS+ On vs CS- Off, p=0.32; CS+ On vs CS- On, p=1.0; CS- On vs CS- Off, p=0.013. **l.** Average power of pip-evoked theta (6-10 Hz) signal in the BLA during the CS+ (red) and CS- (blue) in SOM-cre mice expressing eYFP (left) or eArch (right) in the presence or absence of light. Light has no effect on pip-evoked theta power. **eYFP**, n=7, rm-ANOVA: CS: F_1,18_=7.12, p=0.02; Light: F_1,18_=0.03, p=0.56; CS x Light interaction: F_1,18_=0.05, p=0.83; **eArch**, n=6, rm-ANOVA: CS: F_1,15_=6.89, p=0.02; Light: F_1,15_=0.19, p=0.67; CS x Light interaction: F_1,15_=0.65, p=0.43. **m.** Average pip-evoked theta (6-10 Hz) phase consistency in the PL during the CS+ (red) and CS- (blue) in SOM-cre mice expressing eYFP (left) or eArch (right) in the presence or absence of light. As in the BLA (**k**), silencing of BLA SOM+ INs leads to a disruption of the normal pattern of theta resetting, though no significant change could be demonstrated for the CS+ or CS- alone. Astericks for eYFP indicate the main effect of CS, while notations for eArch reflect significant Bonferroni post-hoc test results. **eYFP**, n=7, rm-ANOVA: CS: F_1,18_=8.07, p=0.011; Light: F_1,18_=0.72, p = 0.41; CS x Light interaction: F_1,18_=1.96, p=0.18. **eArch**, n=6, rm-ANOVA: CS: F_1,15_=13.3, p=0.002; Light: F_1,15_=0, p=0.96; CS x Light interaction: F_1,15_=5.66, p=0.031. Bonferroni post-hoc tests: CS+ Off vs CS+ On, p=0.22; CS+ Off vs CS- Off, p=0.0031; CS+ Off vs CS- On, p=0.44; CS+ On vs CS- Off, p=0.28; CS+ On vs CS- On, p=1.0; CS- On vs CS- Off, p=1.0. **n.** Average power of pip-evoked theta (6-10 Hz) signal in the PL during the CS+ (red) and CS- (blue) in SOM-cre mice expressing eYFP (left) or eArch (right) in the presence or absence of light. Light had no effect on theta power. **eYFP**, n=7, rm-ANOVA: CS: F_1,18_=30.71, p<0.001; Light: F_1,18_=0.31, p = 0.59; CS x Light interaction: F_1,18_=0.26, p=0.62. **eArch**, n=6, rm-ANOVA: CS: F_1,15_=18.85, p<0.001; Light: F_1,15_=0.12, p=0.74; CS x Light interaction: F_1,15_=0.27, p=0.61. *p<0.05, **p<0.01, ***p<0.001. All data plotted as mean +/- SEM.

During DFC retrieval, light selectively enhanced freezing to the CS- in mice expressing Arch in SOM (Figure 2e1) but not in PV neurons (Figure 2e2). This is a particularly strong effect, given that C57/B6 mice are excellent discriminators (Supplementary Figure 11). Light decreased the difference in CS+ and CS-freezing by about 50% for SOM-Arch (Figure 2f1) but not for PV-Arch mice (Figure 2f2), consistent with a substantial, although incomplete, impairment of discrimination. eYFP-expressing controls did not show these effects. Given a possible effect of light in controls, we confirmed that the light-induced reduction in differential freezing to the CSs was significantly different from controls for SOM-Arch animals, but not PV-Arch animals (Supplementary Figure 5).

Given that SOM INs in the BLA mediate distal dendritic inhibition of PNs [13] we tested whether BLA SOM cells gate incoming sensory information, which reliably induces fear-related synchronous activity such as theta oscillation reset in the BLA-PFC circuit[11] [12]. We recorded local field potentials (LFPs) in the BLA and PL in the presence and absence of BLA SOM IN optogenetic silencing. We quantified CS-evoked theta (6-10 Hz) phase reset (Figure 2g, i-k,m, Supplementary Figure 6), theta power (Figure 2h, l, n), and theta coherence (Supplementary Figure 7-8) in the BLA and PL. In the absence of any manipulation, the CS+ but not the CS-evoked phase reset of the ongoing theta oscillations (Figure 2g, k; quantified by phase consistency, Supplementary Figure 6). Cue-evoked reset is a signature of fear learning, as it was not present during habituation (Supplementary Figure 6). In SOM-eArch mice (n=6), SOM inhibition resulted in theta phase reset induction by the CS- with similar temporal dynamics to the CS+, without a significant effect on CS+ induced oscillations (Figure 2i-k). Interestingly, we found that BLA SOM inhibition also equalized the CS+ and CS-theta reset strength of the PL LFP (Figure 2m), although, unlike in the BLA, there was no statistically significant effect detected for CS+ or CS-alone. There was no significant effect in SOM-eYFP animals (n=7, Figure 2k, m). Notably, the SOM silencing effect on theta reset was highly specific, as light did not change BLA or PL theta power (Figure 2l, n), BLA-PL theta coherence or PL-to-BLA direction of information transfer during the CS- (Supplementary Figure 7-8). Thus, BLA SOM activity selectively gates incoming sensory input during CS-presentations, specifically preventing theta reset in the BLA and PL, thereby inhibiting recruitment of circuits mediating defensive behavior.

Given that the PL is important for acquisition of discrimination learning[14], and PL communication with the BLA actively contributes to fear discrimination retrieval [12, 15] [11, 16] [17], we hypothesized that the PL may modulate SOM activity during DFC retrieval. First, we verified that PL activity is necessary for fear discrimination retrieval. We trained mice on DFC and infused saline and muscimol in the PL on different days of retrieval (counterbalanced). Mice froze significantly more to the CS+ than the CS- when saline was infused, whereas muscimol induced fear generalization to the CS-, without an effect on CS+ freezing (Figure 3a-b). In a subset of the mice shown in Figure 1, we imaged SOM INs during DFC retrieval before and after infusion of saline or muscimol in the PL (Figure 3c). PL silencing with muscimol selectively impaired the CS-pattern of calcium activation in SOM cells in all animals (Figure 3d-e, Supplementary Figures 9-10, n=117 neurons, n=3 mice), whereas saline in the PL had no effect on cue processing in SOM cells (Supplementary Figures 9-10, n=127 cells, n=3 mice). PL silencing selectively diminished SOM responses to the CS-but not to the CS+ (Figure 3d-e, Supplementary Figure 9). Whereas SOM responsiveness was significantly stronger to the CS-than to the CS+ prior to muscimol in the PL, there was a slightly higher response to the CS+ than CS-after PL silencing (Figure 3d-e). Accordingly, there was a selective decrease in the percent of CS-responsive cells, with 58% of cells activated significantly by the CS- pre-muscimol in the PL and only 19% post-muscimol (Figure 3f1; Supplementary Figure 10), while saline in the PL had no significant effect on percent of active cells (Figure 3f2). A cell classifier confirmed that encoding remained stable post-saline but not post-muscimol infusion (Figure 3g).

**Figure 3:**
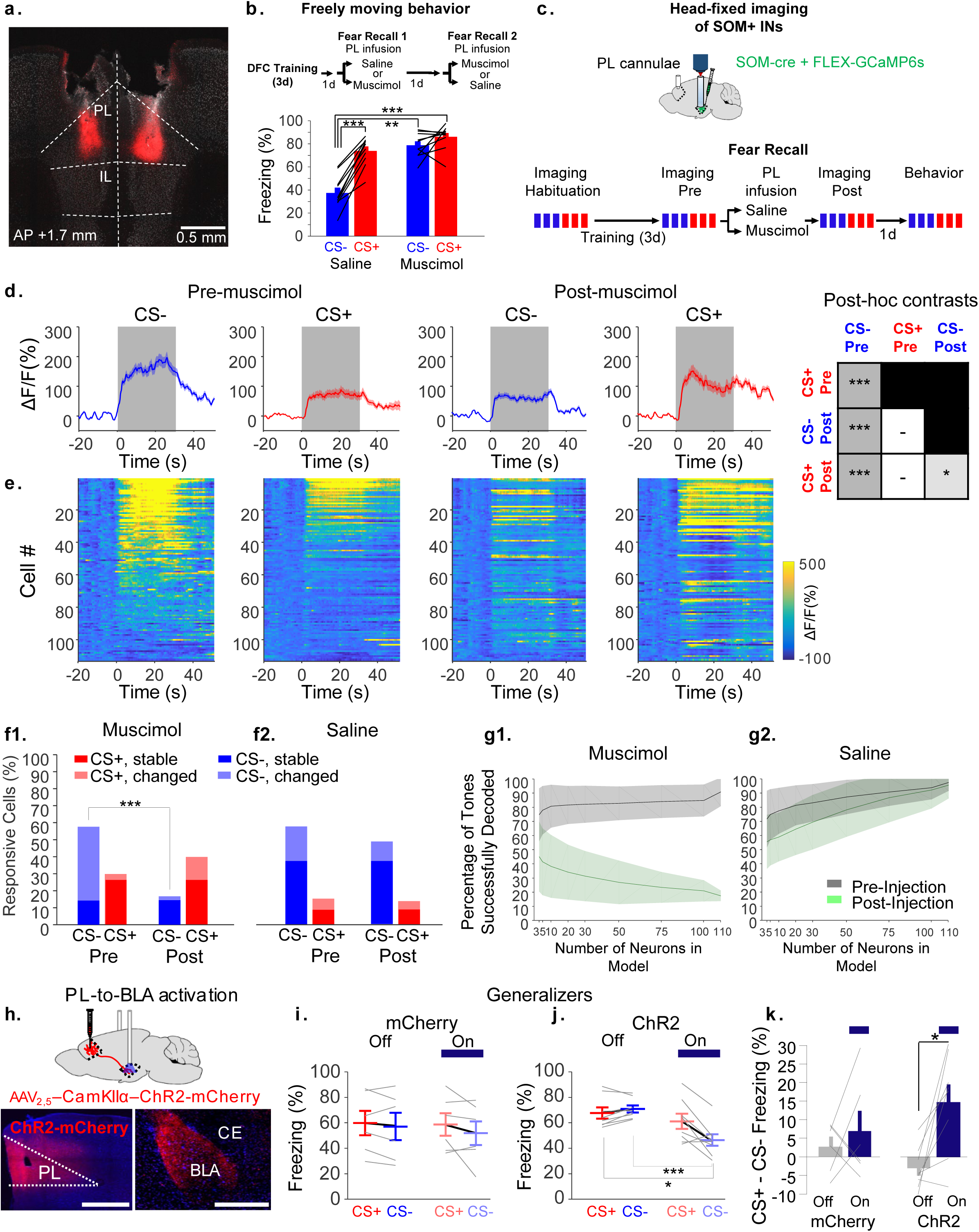
BLA SOM+ activity during the non-threatening stimuli depends on the prelimbic cortex. **a.** Cannulae were implanted above the PL of mice for the infusion of saline or muscimol. DiI of the same volume as used for behavioral experiments was infused through the cannulae prior to making histological sections to visualize the extent of infusions. Histology from a representative mouse is shown. **b.** Mice that were trained on DFC were tested following infusion of saline or muscimol into the PL on subsequent days (order randomized between mice). Muscimol in the PL selectively increased freezing to the CS-, without altering freezing to the CS+. n=10, rmANOVA, CS: F_1,27_=2.23, p =.3755; Drug: F_1,27_=3.34, p=.3185; CS x Drug: F_1,27_=16.15, p<.001. Bonferroni post-hoc tests: CS- Saline vs CS+ Saline, p=2.1 × 10^−5^; CS- Saline vs CS- Muscimol, p=0.002; CS- Saline vs CS+ Muscimol, p=4.6 × 10^−5^; CS+ Saline vs CS- Muscimol, p=1.0; CS+ Saline vs CS+ Muscimol, p=0.083; CS- Muscimol vs CS+ Muscimol, p=0.93. **c.** Experimental design for assessing the effect of PL silencing on SOM+ IN activity during differential fear retrieval. Mice expressing GCaMP6s in the BLA were implanted with cannulae over the PL for the infusion of muscimol or saline, as well as GRIN lenses over the BLA to image SOM IN activity. These mice are a subset of the mice in Figure 1, and they were imaged before and after infusion. **d.** Average CS+ (red) and CS- (blue) evoked calcium signals across all cells in mice infused with muscimol (n=117 cells from n=3 mice) pre- and post-infusion. rm-ANOVA, CS: F_1,336_=48.46, p<.0001; Session: F_1,336_=8.68, p=0.003; CS x session: F_1,336_=57.54, p<.0001. Bonferroni post-hoc test results are summarized on the right. White cells indicate no significant difference and gray cells indicate significant differences. All pairwise contrasts and corresponding results from saline-infused animals can be found in Supplementary Figure 9. **e.** Heat maps plotting individual cell responses, corresponding with average traces in **d**, sorted by the magnitude of the average stimulus-evoked response (in all heat maps). **f1**,**2.** The percentage of cells significantly active during CS- (blue) or CS+ (red) pre- and post-infusion of muscimol (**f1**, n=117 cells in n=3 mice) or saline (**f2**, n=127 cells in n=3 mice). Cells are further subdivided as cells that were significantly active to the CS- or CS+ both pre- and post-infusion (stable) or only in one imaging epoch (changed). PL muscimol significantly decreased the percentage of cells that were CS- responsive (Bonferroni-corrected p<.001, McNemar’s test) but not CS+ responsive (Bonferroni-corrected p=0.32). Saline had no effect on the percentage of cells responsive to CS+ or CS- (Bonferroni-corrected p-values: CS-, p=.32; CS+, p=1.0). **g1**,**2.** Percentage of tones successfully decoded by neural models trained on pre-infusion data, for predicting tone identities pre-infusion (gray) and post-infusion (green) of muscimol (**g1**) or saline (**g2**). Data is plotted as average performance of models composed of different numbers of neurons, with 95% confidence intervals, subsampled from the entire population of recorded cells. Infusion of saline in the PL had minimal effect on the encoding properties of neurons, while muscimol completely altered neural encoding, such that the decoder actually performed below chance levels, consistent with the majority of cells changing their encoding identity (see Supplementary Figure 10 for further exploration of the nature of this change). **h.** Mice were injected with viruses expressing ChR2-mCherry or mCherry in principal cells of the prelimbic cortex. Top, schematic for activation of PL terminals in the BLA. Bottom left, expression of ChR2-mCherry virus in the PL. Bottom right, ChR2-expressing PL terminals in the BLA. Scale bars, 500 microns. **i.** There was no effect of light on freezing in mCherry-expressing mice. n=6, rm-ANOVA, CS: F_1,15_=2.06, p=0.17; Light: F_1,15_=0.91, p=0.36; Light x cue: F_1,15_=0.39, p=0.54. **j.** There was an effect of light on freezing to the CS- in ChR2-expressing mice. n=8, rm-ANOVA, CS: F_1,21_=3.68, p=0.07, Light: F_1,21_=26.51; p<0.0001, Cue x Light, F_1,21_=8.76, p=0.0075, Post hoc Bonferroni tests: CS+ Off vs CS+ On, p=1.0; CS+ Off vs CS- Off, p=0.967; CS+ Off vs CS- On, p=0.0138; CS- Off vs CS- On, p=0.0054; CS+ On vs CS- On, p=0.11; CS- Off vs CS+ on; p=0.45. **k.** The difference in freezing between the CS+ and CS- is plotted for mCherry and ChR2 animals when fiber optic illumination was On or Off. PL terminal illumination led to an increase in discrimination in ChR2-expressing mice (Bonferroni-corrected paired t-test, p=0.017) but not control mice (Bonferroni-corrected paired t-test, p=1.0). *p<0.05, **p<0.01, ***p<0.001. Unless specified, data plotted as mean +/- SEM.

Using mice that typically do not discriminate well during DFC retrieval (129SvEv/Tac mice; Supplementary Figure 11), we investigated whether enhancement of PL input to the amygdala was sufficient to rescue discrimination. We expressed ChR2 (n=8) in the PL and stimulated terminals in the BLA with 6 Hz oscillatory illumination, which selectively decreased CS-freezing, resulting in a significant increase in discrimination score (n=6; Figure 3i-j, Supplementary Figure 12; no effect in controls).

In conclusion, we demonstrate a PL-dependent activation of BLA SOM INs that prevents fear generalization via selective gating of sensory input. During neutral stimuli, BLA SOM INs are active, blocking theta reset of both BLA and PL theta oscillations, consistent with evidence that aversion-related prefrontal theta resets originate in the BLA[17]. When shaping BLA output, the PL could recruit SOM INs indirectly via its numerous contacts with excitatory neurons [18-20] and by direct input. Inhibiting SOM activity decreases inhibition at BLA dendrites, thereby strengthening theta reset, and creating a window for enhanced PL-to-BLA excitatory transmission and plasticity [21], which is likely to induce fear generalization. Notably, PL inactivation blocks BLA SOM INs recruitment during the CS-, and thereby diminishes discrimination. However, theta frequency stimulation of PL inputs to the BLA rescues discrimination, demonstrating an essential role for PL inputs to the BLA during discrimination learning.

## Acknowledgements

We thank Phebe Warren and Estelle Hofgaertner for providing additional technical assistance with histology and behavioral experiments during the revision process.

## Author Contributions

J.M.S. wrote custom software for the analysis. P.K.O-N. performed GRIN lens implantation surgeries, calcium imaging, cannulation surgeries and muscimol inactivation experiments. J.M.S., S.G., L.D., W.H., T.J.S., and E.L. performed surgeries and experiments for *in vivo* optogenetics, electrophysiology, immunohistochemistry and behavior. I.N., S.G., L.D., M.L. provided assistance with behavioral experiments, analysis, imaging and cell counting. S.S.B. and R.T.R. provided assistance with rabies viral tracing. C.D.S. provided assistance with data collection. E.L, J.A.G., J.M.S., designed the experiments, analyzed and interpreted the data, and wrote the manuscript.

## Data Availability

The datasets generated during and analyzed during the current study are available from the corresponding author on reasonable request.

---

**Supplementary Figure 1.**
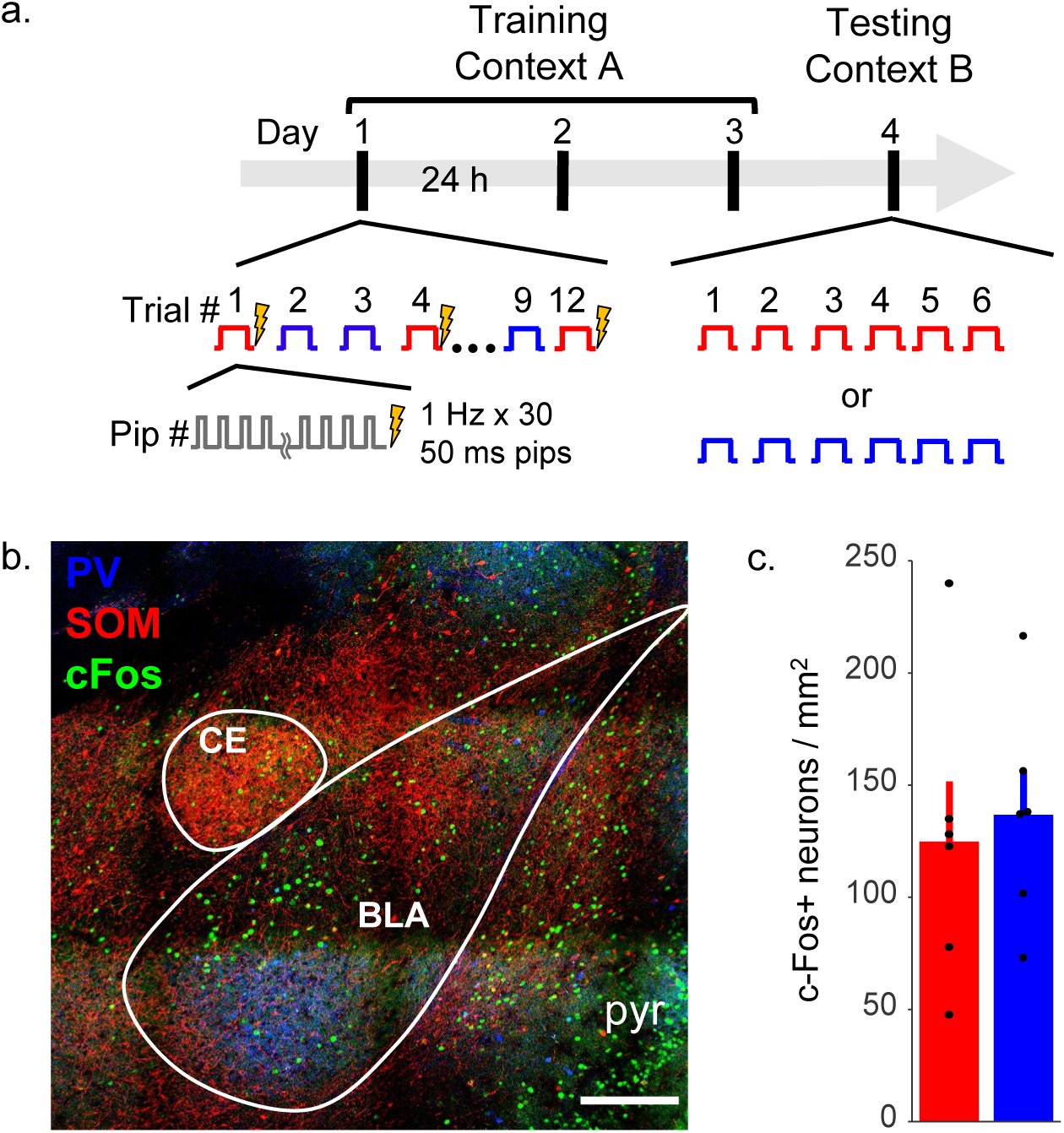
Differential fear learning was assessed by immunohistochemical methods. **a.** DFC training and single CS retrieval paradigm. **b**. An example of c-Fos, PV, and SOM staining after retrieval of the CS-. **c.** The density of cFos+ cells in the BLA does not differ during CS+ and CS- retrieval (n=6 CS+ and n=6 CS-, unpaired t-test, p = .72).

**Supplementary Figure 2.**
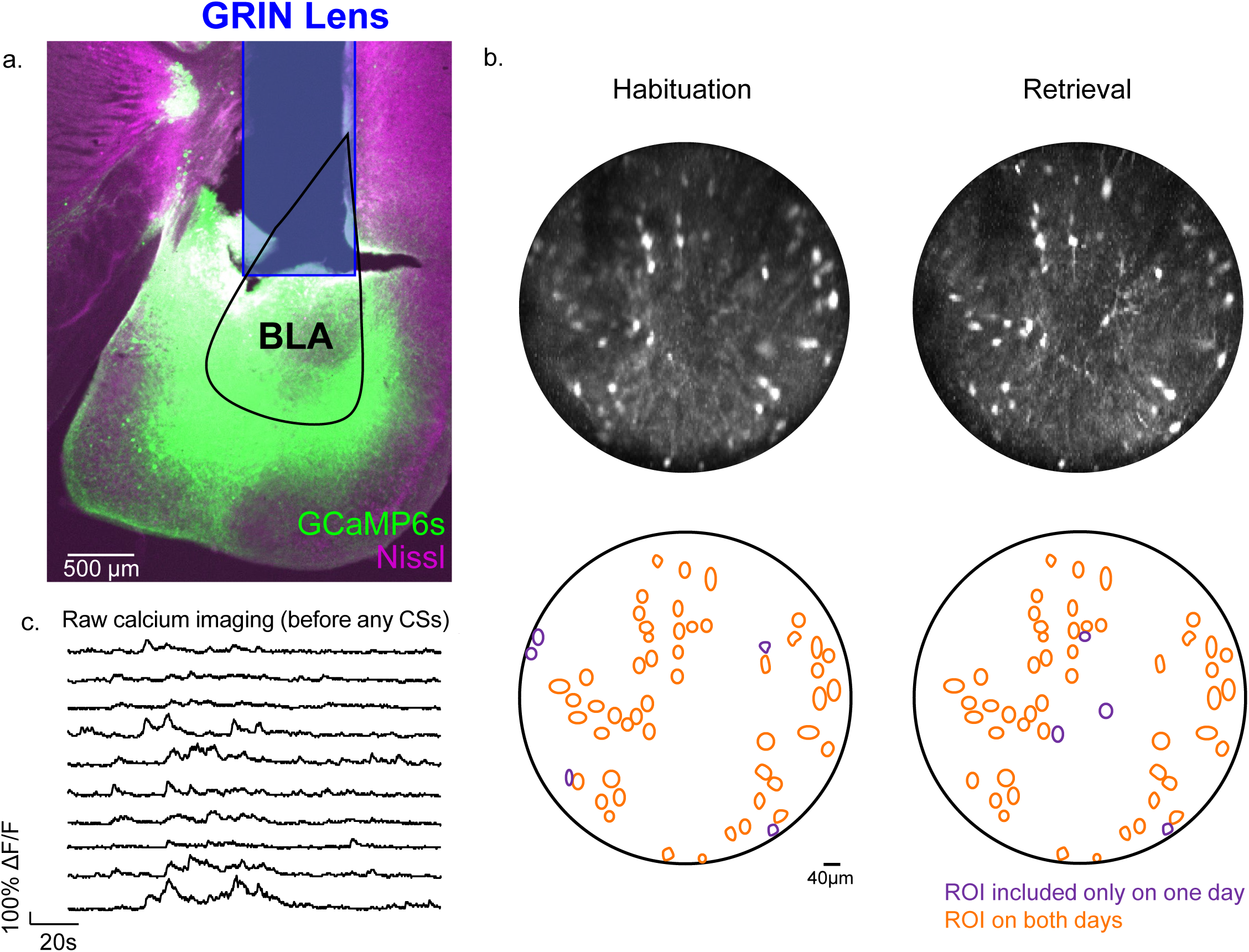
Calcium imaging captured SOM+ IN activity during habituation and retrieval of fear discrimination retrieval. **a.** Example placement of lens (imputed location indicated by blue box) in the BLA of a SOM-cre animal injected with FLEX-GCaMP6s virus. **b.** Top, example projection image of full field of view from one animal during habituation (left) and fear discrimination retrieval (right) after image registration (alignment). Note that the image from habituation is less sharp due to the procedure of registering the video to retrieval day, such that fine features are not as well preserved, without an effect on the activity captured from cell bodies. Bottom, ROIs were mostly stable between habituation and retrieval. A minority of cells were included for only one day (purple), while most cells were imaged on both days (orange). **c**. Examples of raw traces recorded from SOM+ neurons in the BLA of one representative animal during the time period prior to tones being played on retrieval day. In this case, ΔF/F is calculated relative to average of the full recording session.

**Supplementary Figure 3.**
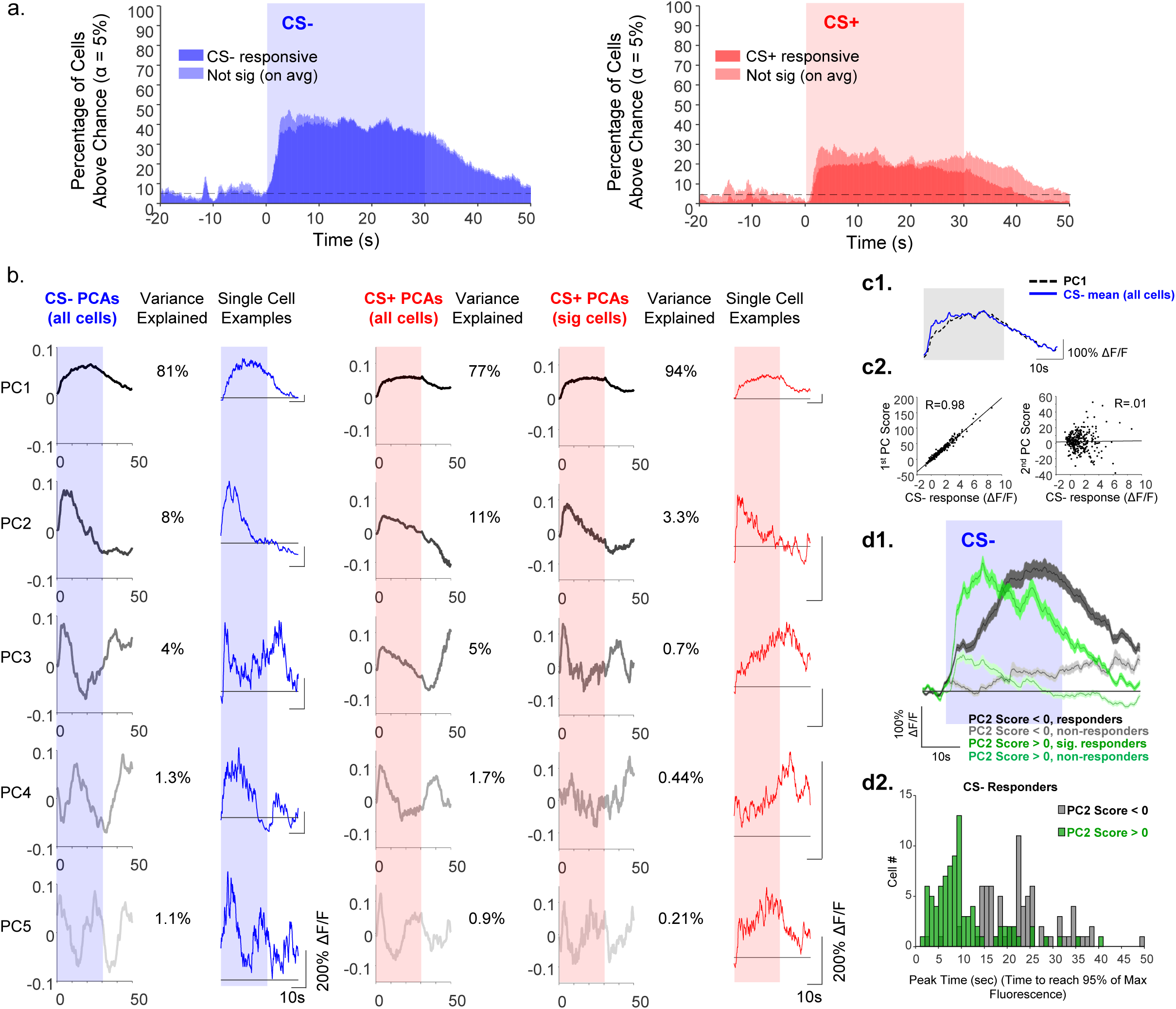
SOM+ INs are homogeneously activated by CSs, with stronger and more consistent response to the CS-. **a.** SOM+ INs (n=306 from 7 mice) were first classified as significantly responsive to the CS- (54% of cells) or CS+ (22% of cells) across the entire 30s tone (compared to that expected by chance). All cells were then assessed for a significant activation on a moment by moment basis. More cells were activated to the CS- (left) than the CS+ (right), but with the same time course, such that activation above chance level could be seen by ∼2 seconds after CS onset. CS evoked activity decreased to baseline over the 20 sec following tone offset. This analysis does not differentiate between cells that were only activated by one CS and those activated by both. **b.** Principal Component Analysis of calcium activity in SOM+ cells during retrieval of CS- and CS+. The first principal component (PC1) explains 81% of the variability of the CS- evoked response among all cells (blue), and 77% of the variability of the CS+ evoked response (red). Principle components are plotted from black (PC1) to light gray (PC5), with quickly diminishing explanatory power. Principal components revealed different patterns of activation that were reflected in individual cells (blue and red traces). When analyzing the response of all cells to the CS- and CS+, principal components were similar but with considerable differences of PC2-5. After restricting the analysis to only CS+ responding cells, PCA components closely matched what was found for the CS-. Note different vertical scaling for different cells. Individuals cell examples are not the same cells for CS+ and CS-. **c1.** Calcium activity described by PC1 (hatched black) closely matches the mean SOM+ response to the CS- (blue). **c2.** (Left) PC1 score is significantly correlated with the CS-response (left, R=0.98, p < .001), whereas (Right) PC2 Score is not significantly correlated with the CS- response (R=.01, p > .05). **d1.** PC2 captures different latencies of response. CS- responsive cells with positive PC2 scores (dark green) have a faster onset and offset dynamic than those with negative PC2 scores (dark grey). Cells with non-statistically significant response to the CS- are included for comparison (light hues). **d2.** Overallping histograms of peak fluorescence time in CS- responsive cells reveals a bimodal distribution, such that cells with positive (green) and negative (gray) PC2 scores have short and long latency responses, respectively.

**Supplementary Figure 4.**
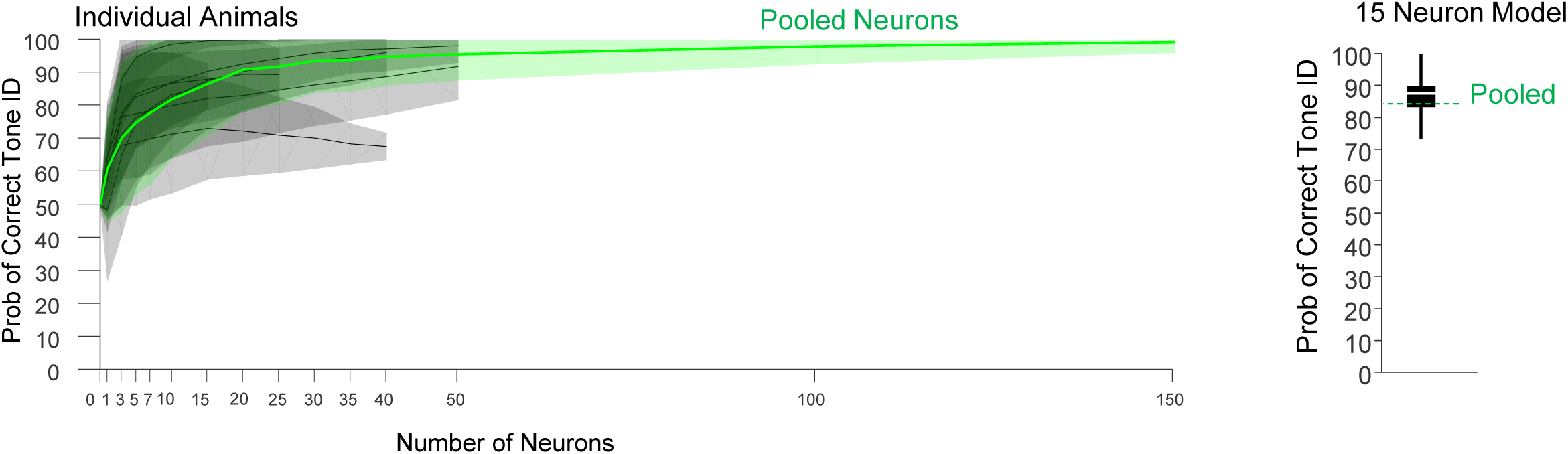
Cell classifier decodes tone identity based on SOM+ activity in each animal. Left panel, Probability of correctly identifying the tone as a CS+ or a CS- based on the number of neurons included in the model for each recorded animal (grey, n=7 mice), and when neurons were pooled (green, n=306 total) from all animals. Due to differences in the number of neurons recorded from each mouse, the curves are of different lengths. Right panel, Box and whisker plot depicting the decoding accuracy in individual animals, using 15 subsampled SOM INs, relative to the pooled average, shown in green. Group median probability of correctly identifying tone identity with 15 subsampled SOM+ neurons indicated by white line, 25% and 75% quartiles across animals shown by black box, and maximum an minimum values marked by black whiskers. The decoding reliability was different from chance (one-sample t-test vs. 50%, p=2.4 × 10^−5^).

**Supplementary Figure 5.**
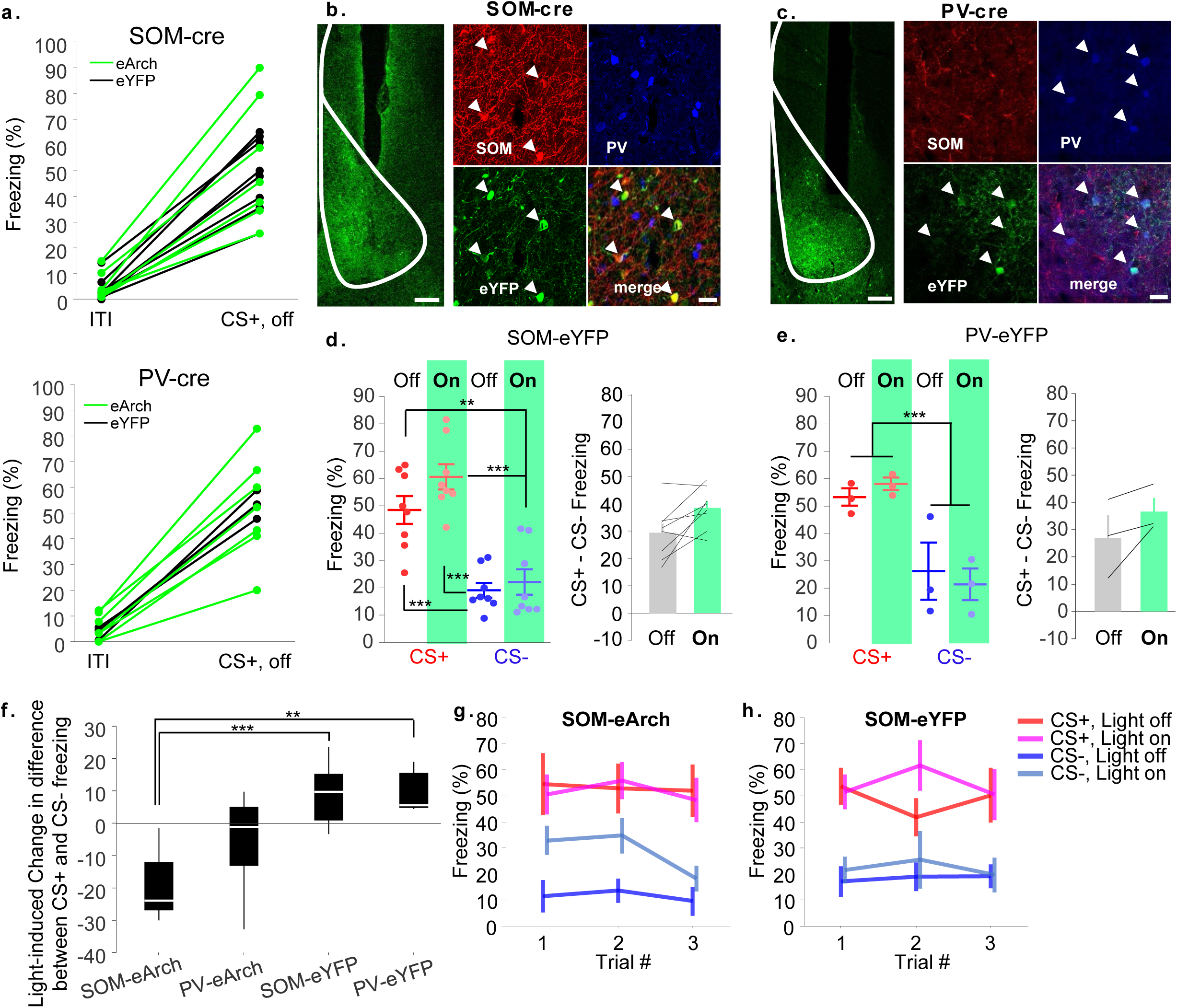
Optogenetic silencing of SOM INs increases CS- freezing without altering CS+ freezing. **a.** All mice increased freezing by over 20% during the CS+ compared to the ITI. Each SOM-cre (top) and PV-cre (bottom) mouse is indicated by each line, color coded by virus (Arch, green vs eYFP, black). ITI freezing, 4.64 +/- 0.9% vs. CS+ off, 51.4 +/-3.5%. **b-c.** Left, Example histology showing optic fiber placement in a mouse expressing FLEX-Arch in SOM-cre (b) or PV-cre (c) mice. Right, example immunohistochemical characterization of cells expressing FLEX-eYFP (green), demonstrating reliable expression in non-overlapping populations of SOM-expressing (red) and PV-expressing (blue) cells. **d-e**. Freezing behavior of control mice expressing eYFP in SOM-cre (d) and PV-cre (e) mice, as in Figure 2. **d.** SOM-cre + FLEX-eYFP, n=8, rm-ANOVA: CS: F_1,21_=217.42, p<.001; light: F_1,21_=10.92, p=.003; CS x light: F_1,21_=3.91, p=.06. Post hoc Bonferroni tests: CS+ Off vs CS+ On, p=.07; CS- Off vs CS- On, p=1.0; CS+ Off vs CS- Off, p=4×10^−5^; CS+ Off vs CS- On, p=.003; CS+ On vs CS- On, p=1.14×10^−5^; CS- Off vs CS+ on, 1×10^−5^. **e.** PV-cre + FLEX-eYFP, n=3, rm-ANOVA: CS: F_1,6_=53.05, p=.0003; light: F_1,6_=0, p=1.0; CS x light: F_1,6_=1.21, p=.31. Bonferroni-corrected paired t-test for Difference in Freezing, Light Off vs On: p=1.0. **f.** Box plot depicting change in CS+ - CS- difference for SOM-eArch (n=7), PV-eArch (n=7), SOM-eYFP (n=8), and PV-eYFP (n=3) animals, which was only significantly different in SOM-eArch animals. One-way ANOVA: F_3,21_=9.29, p=4×10^−4^. Bonferroni post-hoc tests: SOM-eArch vs SOM-eYFP, p=5×10^−4^; SOM-eArch vs PV-eYFP, p=.0076; PV-eArch vs SOM-eYFP, p=0.1360; PV-eArch vs PV-eYFP, p=0.42;PV-eYFP vs SOM-eYFP, p=1.0; PV-eArch vs SOM-eArch, p=.18. **g-h.** Trial by trial freezing to the CS+ and CS- during “Light on” and “Light off” trials in SOM-eArch animals (g), and SOM-eYFP controls (h).

**Supplementary Figure 6.**
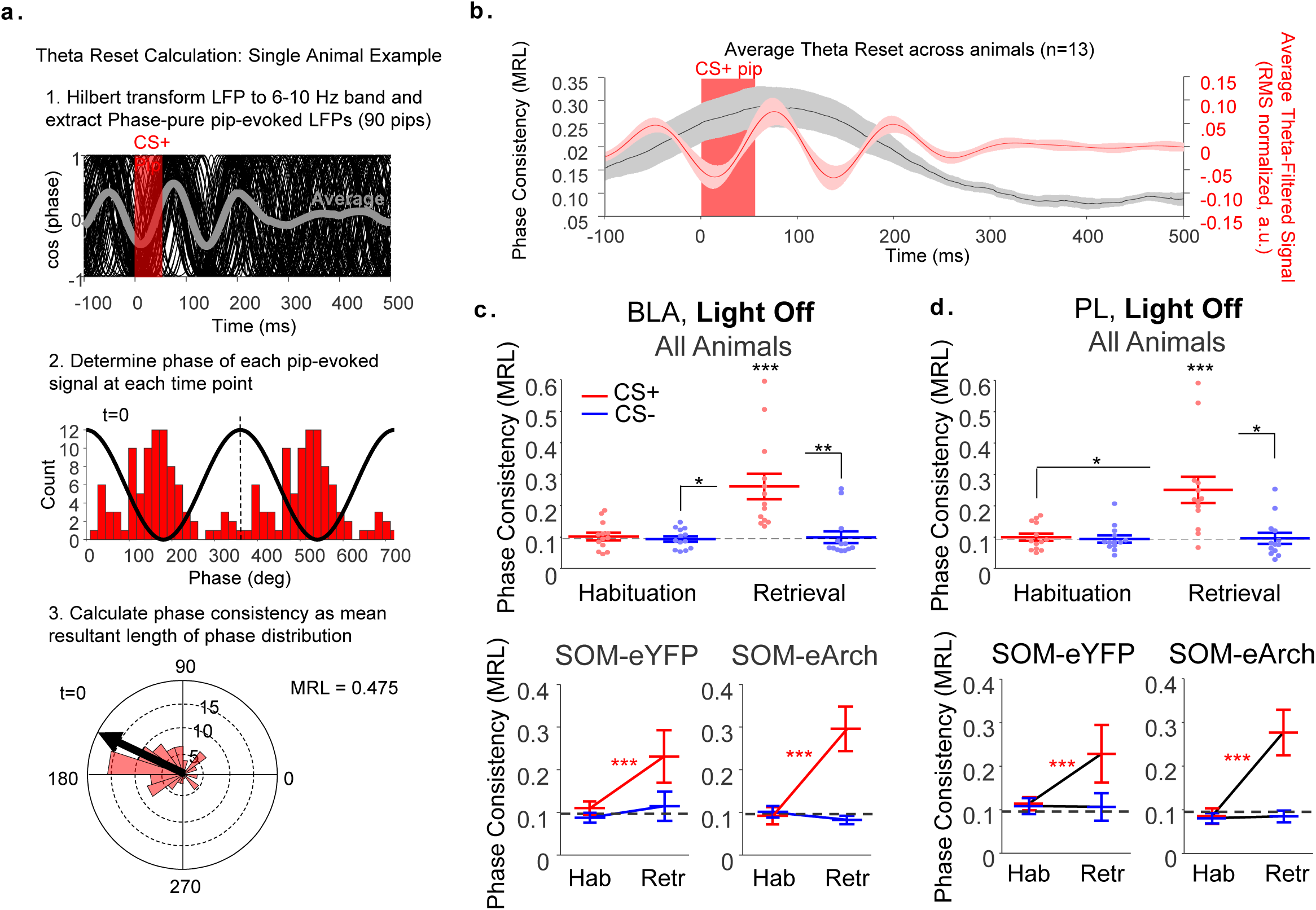
After DFC, the CS+ but not the CS- evokes a reset of the ongoing theta phase. **a.** Theta phase analysis. Steps in the analysis of cue-evoked theta phase consistency (theta reset). **1.** The data is filtered using the Hilert transform for the theta band (6-10 Hz), and phase information is extracted. A one-second filter size was used to ensure accurate phase determination, with the downside of some loss of temporal specificity, such that theta phase resetting is artificially seen before the pip onset. An example of all 90 responses from a single animal to CS+ pips is plotted as cosine of the phase. **2**. A histogram of phases is constructed at each time point relative to CS pip onset (−100 to 500 ms every 0.5 ms). The histogram for an example animal at time 0 (pip onset) is shown. **3.** The phase consistency is calculated at each time point, as a mean resultant length (MRL) vector of the phase distribution (shown in 2). **b.** Average theta phase consistency (gray) and average theta-filtered LFP signal (red) for all mice (n=13) around CS pip onset during Light Off trials. Theta phase consistency peaked shortly after each pip terminated. A trough of theta is induced by CS+ pip onset, with a peak shortly after pip offset. **c-d.** Theta phase reset was quantified for BLA (**c**) and PL (**d**) during light off for all mice (top) and separately for SOM-eYFP and SOM-eArch (bottom). **c.** Theta resetting, as quantified by average phase consistency between 0-200ms after pip onset compared to chance, was elevated to the CS+ during retrieval in the BLA and PL. **BLA** (**c**): rm-ANOVA: n=13 per day (12 mice paired and 1 mouse unpaired on each day due to electrical noise), Day: F_1,35_=3.42, p=.07; CS: F_1,35_=17.75, p=.0002; Day x CS interaction: F_1,35_=9.55, p=.004. Bonferroni post-hoc tests: CS+ Hab vs CS- Hab, p=1.0; CS+ Hab vs CS+ Recall, p=.14; CS+ Hab vs CS- Recall, p=.77; CS- Hab vs CS+ Recall, p =.018; CS- Hab vs CS- Recall, p=1.0; CS+ Recall vs CS- Recall, p=.0064. **PL** (**d**): rm-ANOVA: n=13 per day (12 mice paired and 1 mouse unpaired), Day: F_1,35_=3.71, p=.06; CS: F_1,35_=13.89, p=.0007; Day x CS interaction: F_1,35_=13.69, p=.0007. Bonferroni post-hoc tests: CS+ Hab vs CS- Hab, p=1.0; CS+ Hab vs CS+ Recall, p=.022; CS+ Hab vs CS- Recall, p=1.0; CS- Hab vs CS+ Recall, p =.07; CS- Hab vs CS- Recall, p=1.0; CS+ Recall vs CS- Recall, p=.013. After identifying that the CS+ during recall was different from the other stimuli, we then investigated whether there were any detectable theta reset above chance levels in each condition. During habituation (prior to fear conditioning), neither the CS+ nor the CS- elicits a theta reset in either the BLA (**c**, n=13) or PL (**d**, n=13), (p>.05, one-sample t-tests, log [MRL] vs log [chance probability, .0935]). During fear retrieval (after conditioning), theta reset is elicited by the CS+ (BLA, and PL, p<.0001, one-sample t-test, log [MRL] vs log [.0935]) but not the CS- (BLA and PL, p>.05, one-sample t-test). The significant increase in CS+ induced theta resetting without a change in CS- induced theta resetting is seen for both SOM-eYFP and SOM-eArch animals when the light is off and SOM+ INs are active (p<.001, paired t-tests, log[MRL]). *** p <.001. Together with the main findings in Figure 2, SOM-eArch and SOM-eYFP demonstrate the same patterns of theta resetting in the absence of light, whereas when the light is turned on, SOM+ IN silencing in SOM-eArch animals allows for CS- induced theta resetting, which otherwise does not occur, neither before nor after DFC.

**Supplementary Figure 7.**
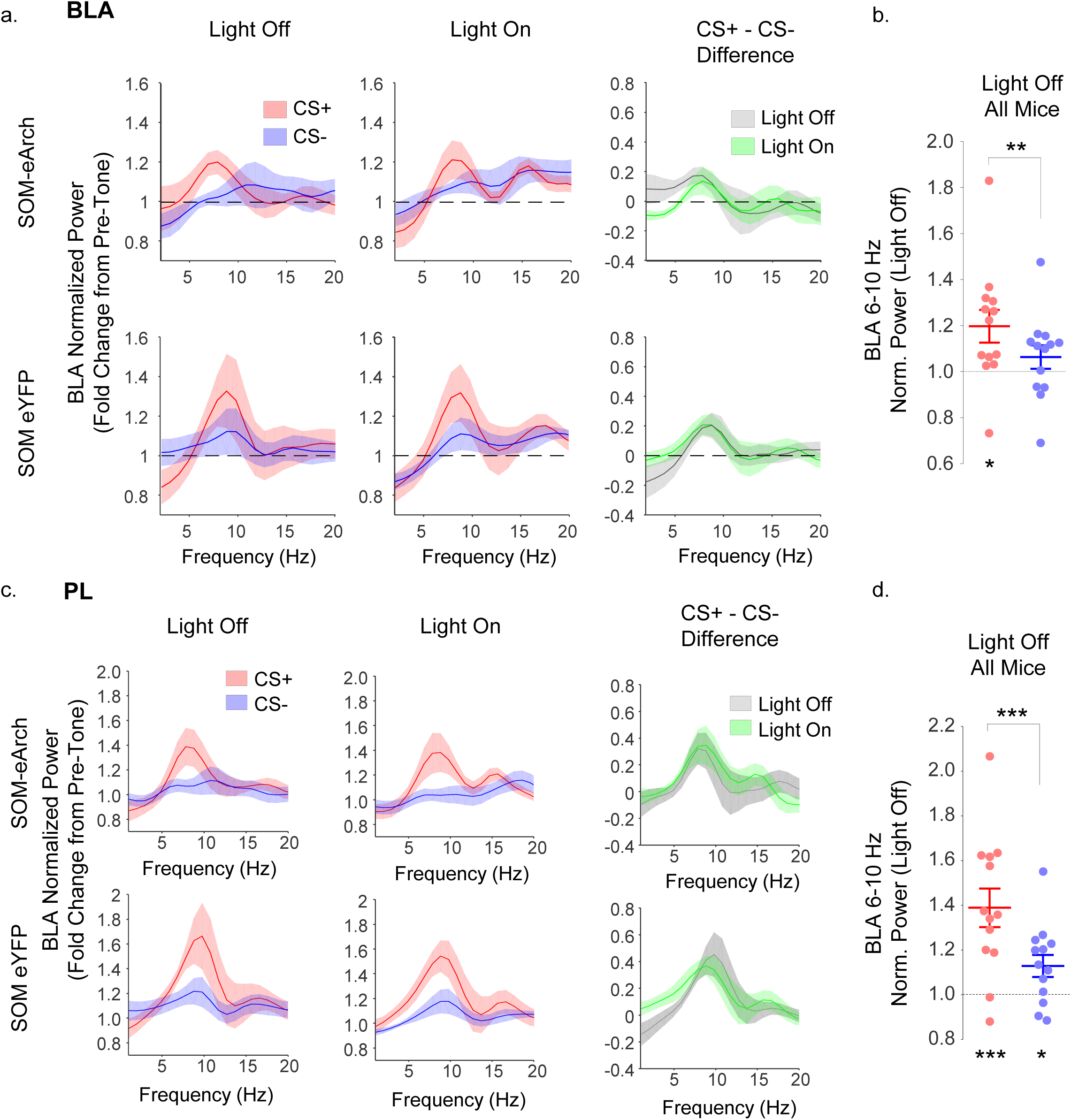
Inhibition of SOM+ cells in the BLA during DFC retrieval does not alter theta power in the BLA or PL. **a.** Normalized power spectra (fold change from pretone) in the eArch (top, n=6) and eYFP (bottom, n=7) groups during Light Off (left), and Light On (middle) trials. The CS+ continues to evoke higher theta power than the CS- in the Light On trials, when SOM+ cells are inhibited (right panel).Broken line, chance levels. **b.** Group data for all mice (n=13) during Light Off trials. CS+ evoked BLA theta power is significantly above pretone (p<.05, one-sample t-test for log[normalized power] vs 0), and above CS- evoked theta power (p<.01, paired t-test for log [norm. power]). **c.** CS evoked power changes from pretone levels in the PL during Light Off trials (left panel) and Light On trials (middle panel) show that the PL theta power is higher during the CS+ than the CS- (subtraction, right panel) in both SOM-eArch (top, n=6) and SOM-eYFP (bottom, n=7) mice, regardless of light. **d.** Group data for all mice (n=13) during Light Off trials. CS+ evoked PL theta power is significantly above pretone (p<.001, one-sample t-test), and above CS- evoked theta power (p<.001, paired t-test). CS- evoked PL theta power was also significantly above pretone (p<.05, one-sample t-test). * p < .05, ** p < .01, *** p <.001

**Supplementary Figure 8.**
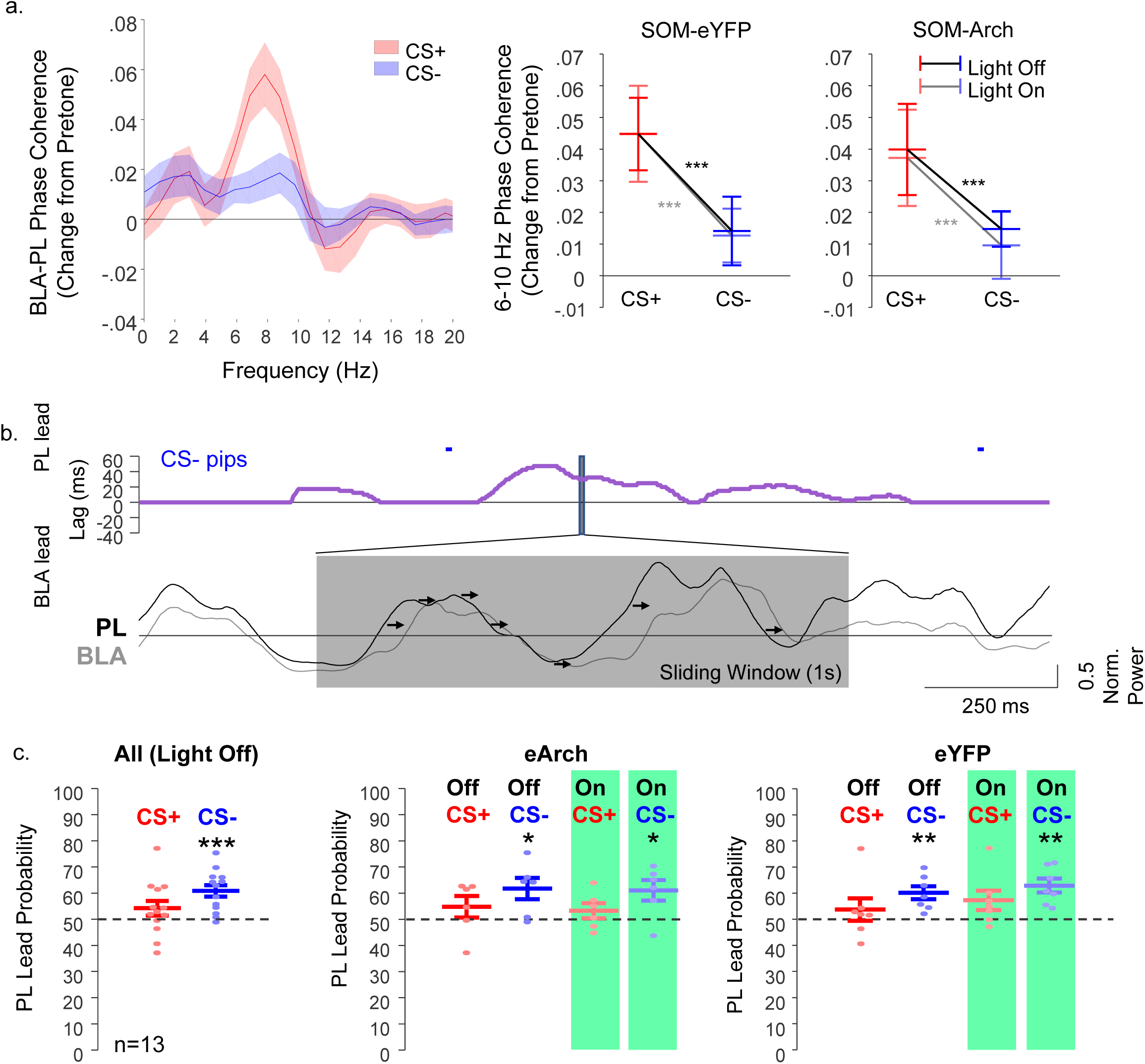
Inhibition of SOM+ cells in the BLA during DFC retrieval does not alter theta coherence or direction of communication between PL and BLA. **a.** Average normalized BLA-PL phase coherence is higher during the CS+ than the CS- during light off trials (left panel). Fiber optic illumination does not affect higher coherence in the CS+ than the CS- in either the SOM-eYFP (middle panel, rmANOVA, n=7, CS: F_1,18_=27.63, p<.001; Light: F_1,18_=.01,p=0.9; CS x Light: F_1,18_=.02, p=0.9) or the SOM-eArch (right panel, rmANOVA, n=6, CS: F_1,15_=24.67, p<.001; Light: F_1,15_=0.54,p=0.47; CS x Light: F_1,15_=.06, p=0.81) groups. *** p < .001, denoting significant effect of CS without effect or interaction with light. No post-hoc testing was performed. **b.** Example of power directionality analysis for simultaneously recorded activity in PL and BLA, showing PL lead over the BLA during the CS- pips (Top panel: CS- pips, blue; directionality, purple line). Lead was calculated in 1s sliding windows by looking at the peak lag of cross-correlation between PL (black) and BLA (grey) average multitaper theta power, and the changes in theta power occurring first in the PL followed by the BLA during the CS-. Arrows show the size of the PL lead in ms. **c.** Left panel, power correlations analyses show the probability of a PL lead during the CS+ (red) and CS- (blue) in all animals (n=13) during Light Off trials. Whereas PL theta oscillations do not lead BLA theta power above chance levels during the CS+ (one sample t-test vs 50%, p=0.17), PL theta changes leads the BLA significantly above chance during the CS- (one sample t-test, p=.003). The PL lead during the CS- is not affected by SOM+ cell inhibition in the BLA of animals in the SOM-eArch group (**middle panel**, one-sample t-test, Light Off CS+, p=.29, CS-, p=.03; Light On CS+, p=.3; CS-, p=.036) or the SOM-eYFP group (**right panel**, one-sample t-test vs 50%, Light Off CS+, p=0.42, CS-, p=.006; Light On CS+, p=0.1; CS-, p=.003). No between group comparison testing was done and no multiple comparisons correction was applied, as difference from 50% was defined a priori as our output metric for this analysis (see methods). Broken line, chance level (50%). * p < .05, ** p < .01, *** p < .001

**Supplementary Figure 9.**
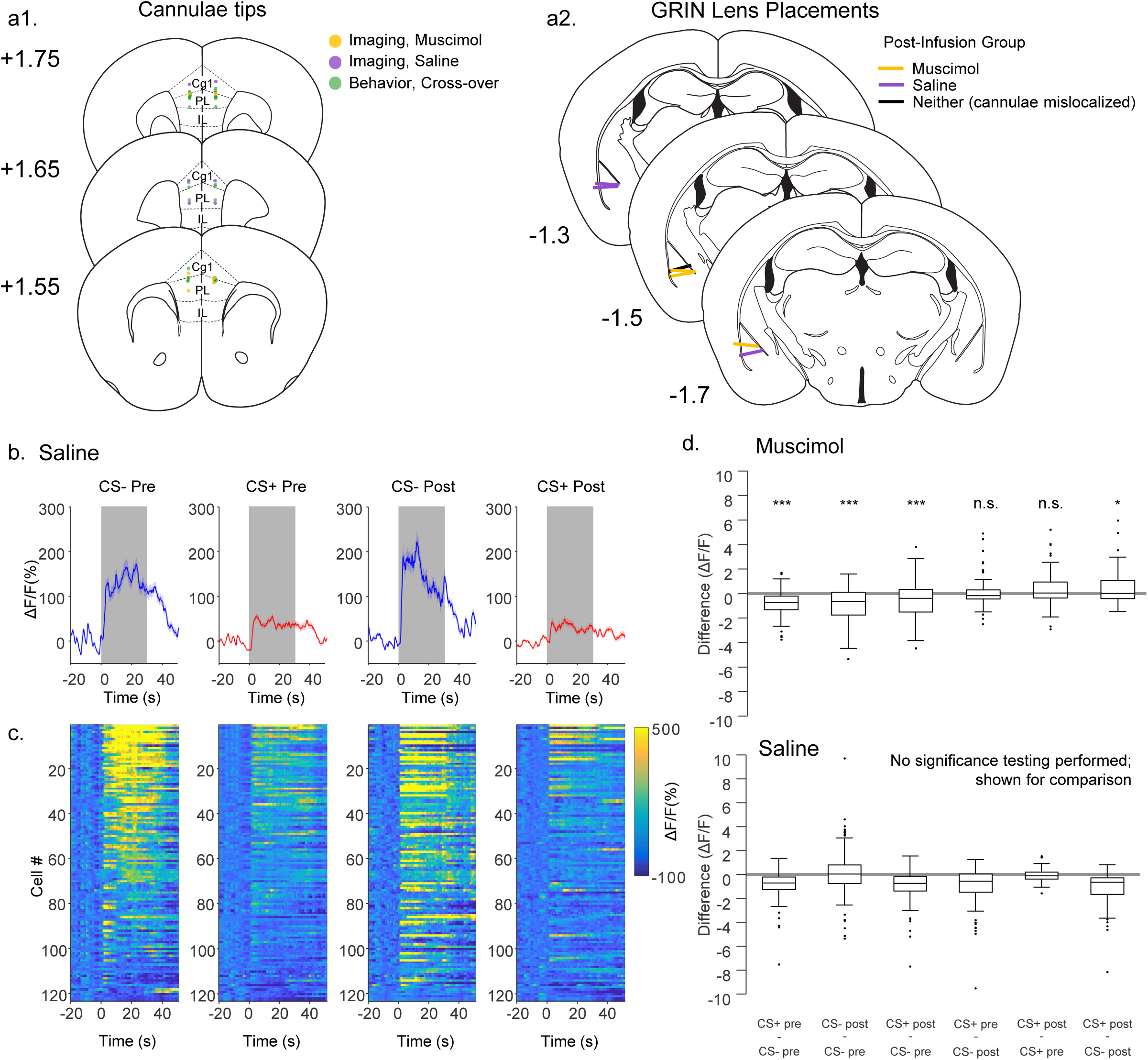
Infusion of muscimol but not saline into the PL disrupts differential BLA SOM activation to the CS-. **a.** Histology for PL manipulation experiments. **a1.** The location of saline/muscimol infusions is included for all animals in **Figure 3** (n=16). Animals used for behavior in the saline-muscimol cross-over design are noted in green, and imaging animals are noted in purple (saline) and orange (muscimol). **a2.** The location of all GRIN lens is noted by lines (n=7 mice, 6 mice that were included in PL infusion experiment (orange, purple) and one (black) that was only included for pre-infusion data in Figure 1 due to PL cannulae that were mislocalized. Numbers denote AP locations. Slices are adapted from Paxinos reference atlas. **b.** Average CS evoked calcium fluorescence of BLA SOM during DFC recall of the CS- (blue) and CS+ (red) before and after saline infusions in the PL. There is no significant change in cue evoked activity. **c.** Heat plots showing activity in all recorded cells that were responsive to the CS- (blue) and CS+ (red) before and after saline infusion in the PL. The same cell is shown in each row, sorted by average response to CS before infusion, as in **Figure 3**. There was no significant effect of saline infusion on the average response to CS- or CS+. rm-ANOVA, n=127 cells from 3 mice, CS: F_1,366_=51.51, p<.0001; Session: F_1,366_=0.34, p=.56; CS x Session: F_1,366_=1.68, p=.196. **d.** Top, boxplots depict all post-hoc contrasts, which were analyzed using post-hoc Bonferroni testing in animals that had muscimol infused in the PL. Results of rm-ANOVA and p-values are reported in legend of Figure 3, though post-hoc significance testing is summarized here. Bottom, equivalent boxplots for saline, but post-hoc Bonferroni testing was not performed due to a lack of significant effect of saline or significant interaction on rm-ANOVA testing. CS denotes CS+ vs CS- and session denotes pre-infusion vs post-infusion categorical variables. Bonferroni post-hoc testing for muscimol group: CS+ pre vs CS- pre, p<.001; CS- post vs CS- pre, p< .001; CS+ post vs CS- pre, p<.001; CS+ pre vs CS- post, p=1.0; CS+ post vs CS+ pre, p=1.0; CS+ post vs CS- post, p<.05* p < .05, *** p < .001

**Supplementary Figure 10.**
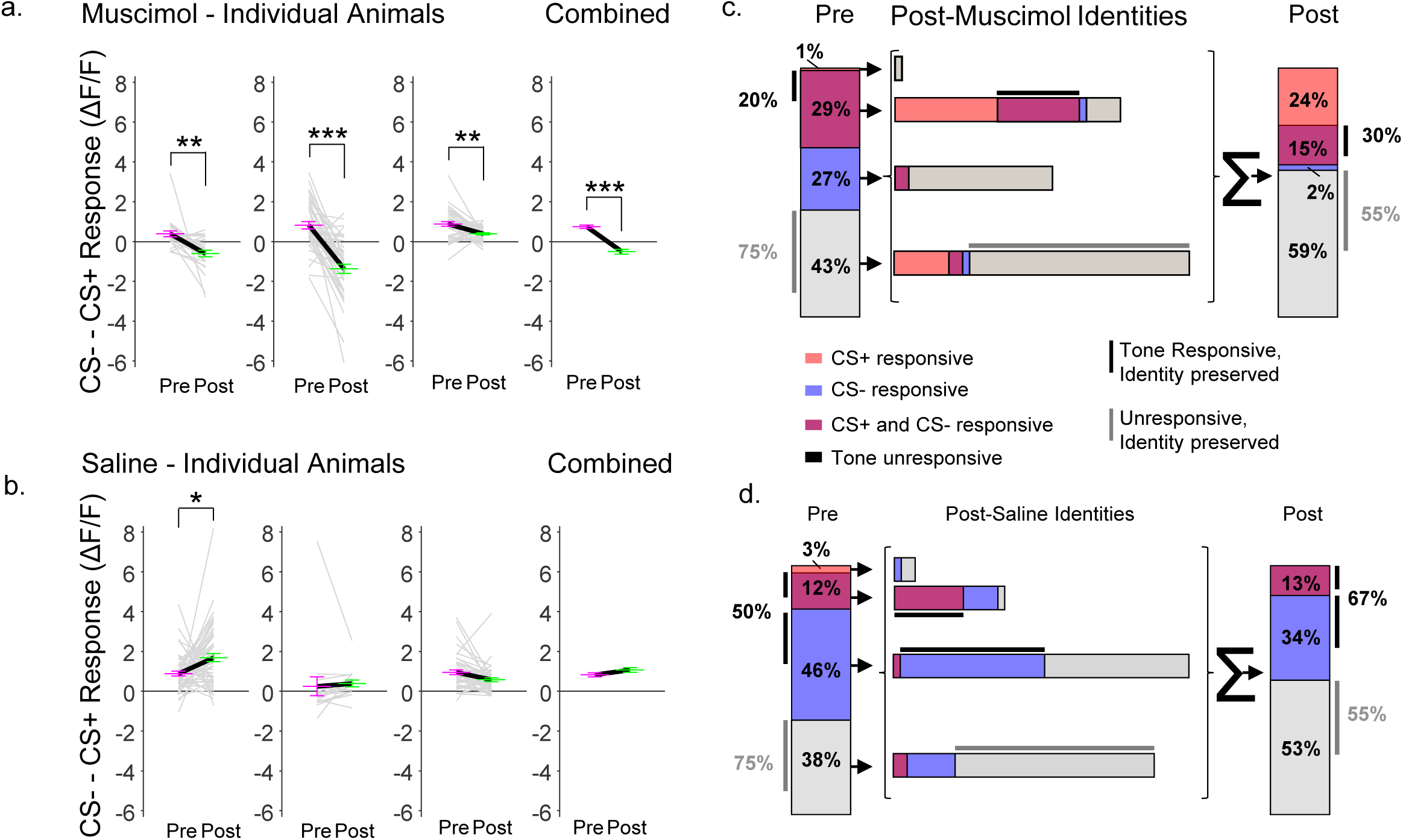
SOM cells in individual animals show a larger response to the CS- than the CS+, and this difference is lost after PL silencing. **a-b.** The difference in ΔF/F between CS- and CS+ is plotted as gray lines for all the cells from each individual animal pre-infusion and post-infusion of muscimol (**a**) or saline (**b**). Average differences are plotted as pink (pre-infusion) and green (post-infusion) for the three mice that received muscimol (a, left three graphs), and the three mice that received saline (b, left three graphs). Combined data is shown in graphs on the right. **a.** Following muscimol infusion, the difference in CS- responding relative to the CS+ is decreased in SOM cells in each individual animal (Bonferroni-corrected paired t-tests with significant change between pre and post for each individual animal and for combined data), whereas the differential response is unchanged in animals that received saline in the PL (**b**), with the exception of one animal in the saline group (left most panel in b) which showed a stronger CS- response relative to the CS+ after saline infusion (Bonferroni-corrected p=.03), but this effect was not seen in the other mice or when data were combined. * p < .05, ** p < .01, *** p <.001 **c-d.** Mapping of responses in all recorded BLA SOM cells pre and post muscimol (**c**) or saline (**d**) infusions in the PL. **Left vertical columns**, breakdown (in percent) of how the recorded cell population responded to CS+ only (red), CS- only (blue), both CS+ and CS- (maroon/purple), or was unresponsive (grey). **The middle, horizontal bars**, show how each segment of the population (e.g. CS+ only responsive) was reorganized in its response patterns after injection of muscimol (**c**) or saline (**d**). For example, the 1% of cells that were CS+ only responsive prior to an injection of muscimol in the PL, became unresponsive to cues after the injection, thereby switching its identity from “CS+ responsive” to “Tone Unresponsive” (**c**, top horizontal bar)., **Right vertical column**, the breakdown (in percent) of how the recorded cell population responded to tones post-infusion. Note that all the responses of a particular type (e.g. CS+ responsive) are summed from the middle, horizontal bars, to account for 100% of the recorded cells. The thin black bars highlight cue-responsive cells that preserved their Tone Identities from before to after the infusion, and the grey bars highlight Unresponsive cells that preserve their Unresponsive Identity from before to after the infusion. Numbers to left and right denote the percentage of pre-infusion identities (left) and post-infusion identities (right) that were the same for tone-responsive (black) and unresponsive (gray) cells. The percentage of cells that retained their tone identity was significantly different between animals infused with muscimol and those infused with saline. No CS- only responsive cells remained responsive to the CS- only following muscimol infusion, and only a minority of cells that responded to both retained a CS- response, in contrast to higher levels of continued response in saline infused animals. Of those that were tone responsive pre-infusion, 20% retained the same tone-responsive identity for muscimol mice compared to 50% in saline mice, χ^2^ (2, N=102) = 13.2, p=.0002; likewise, of those that were tone responsive after infusion, only 30% were of the same identity as before muscimol infusion, compared to 67% as before saline infusion, χ^2^ (2, N=239) = 13.3, p=.0002.

**Supplementary Figure 11.**
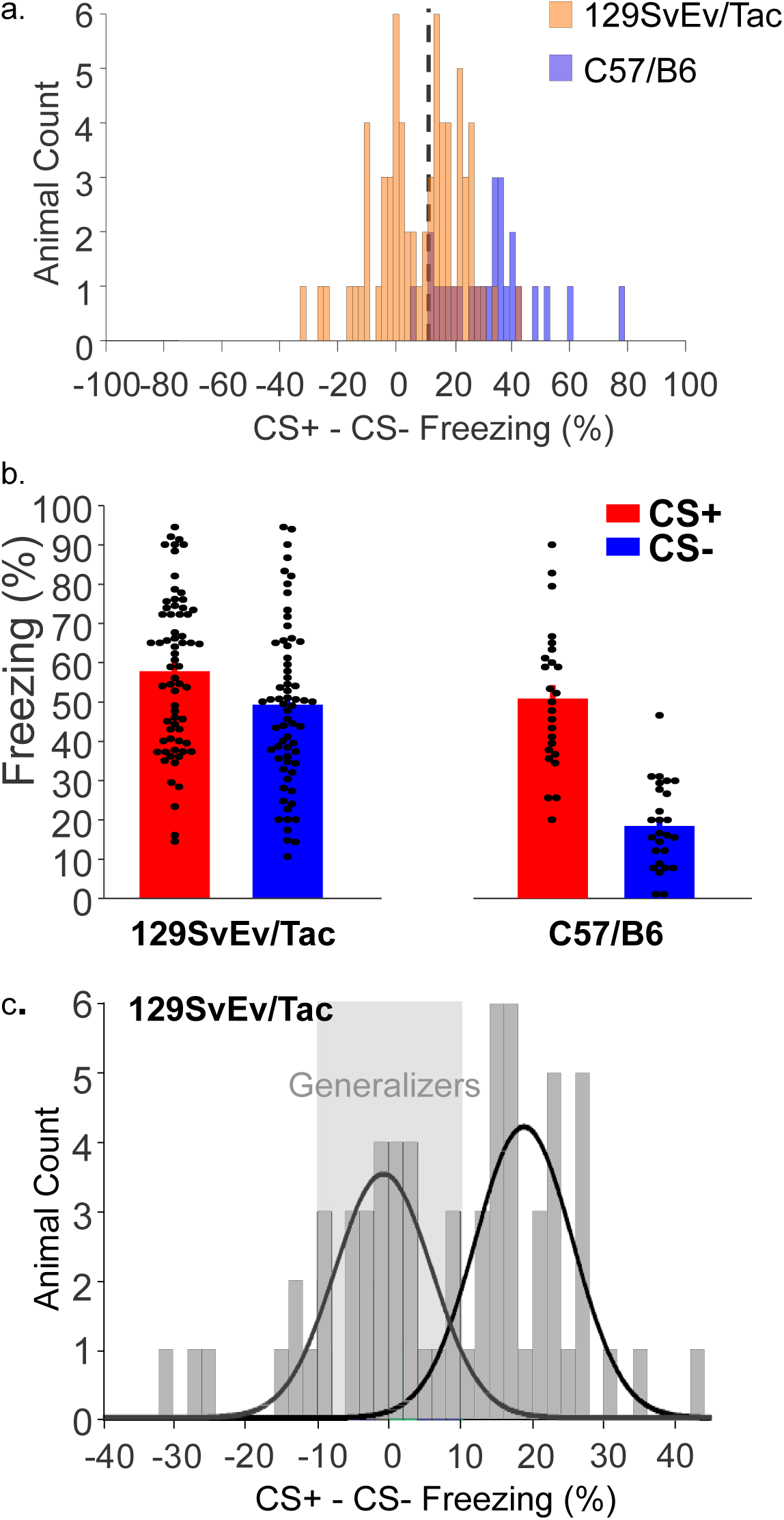
129SvEv/Tac mice show a bimodal pattern of difference between CS+ and CS- freezing, with overall a higher level of generalization compared to C57/56 mice. **a.** Data from 129SvEv/Tac (n=65) and C57/B6 (n=26) mice that underwent differential fear conditioning and then were tested the next day for freezing to the CS+ and CS-. This data includes a combination of mice that were tested in the absence of any manipulation and those used in optogenetic experiments, in which case only Light Off data was used. The difference between CS+ and CS- freezing is shown for all mice, demonstrating that C57/B6 mice are almost all demonstrate a freezing difference greater than 10%, while129SvEv/Tac exhibit a bimodal distribution with a substantial portion of mice with a low difference. Histograms are depicted in an overlapping format. **b.** Data for the same mice is depicted to show CS+ and CS- freezing, showing that in both 129SvEv/Tac and C57/B6 mice, CS+ freezing is on average 50-60%, with divergent distributions of CS- freezing. **c.** 129SvEv/Tac mice were classified as generalizers if the difference between CS+ and CS- freezing was between +/- 10%. Best fit distributions for two overlapping normal distributions is shown by the overlying lines to fit the bimodal distribution, substantiating the basis of these cutoff values. Goodness of fit: R^2^=.812. See methods for further details.

**Supplementary Figure 12.**
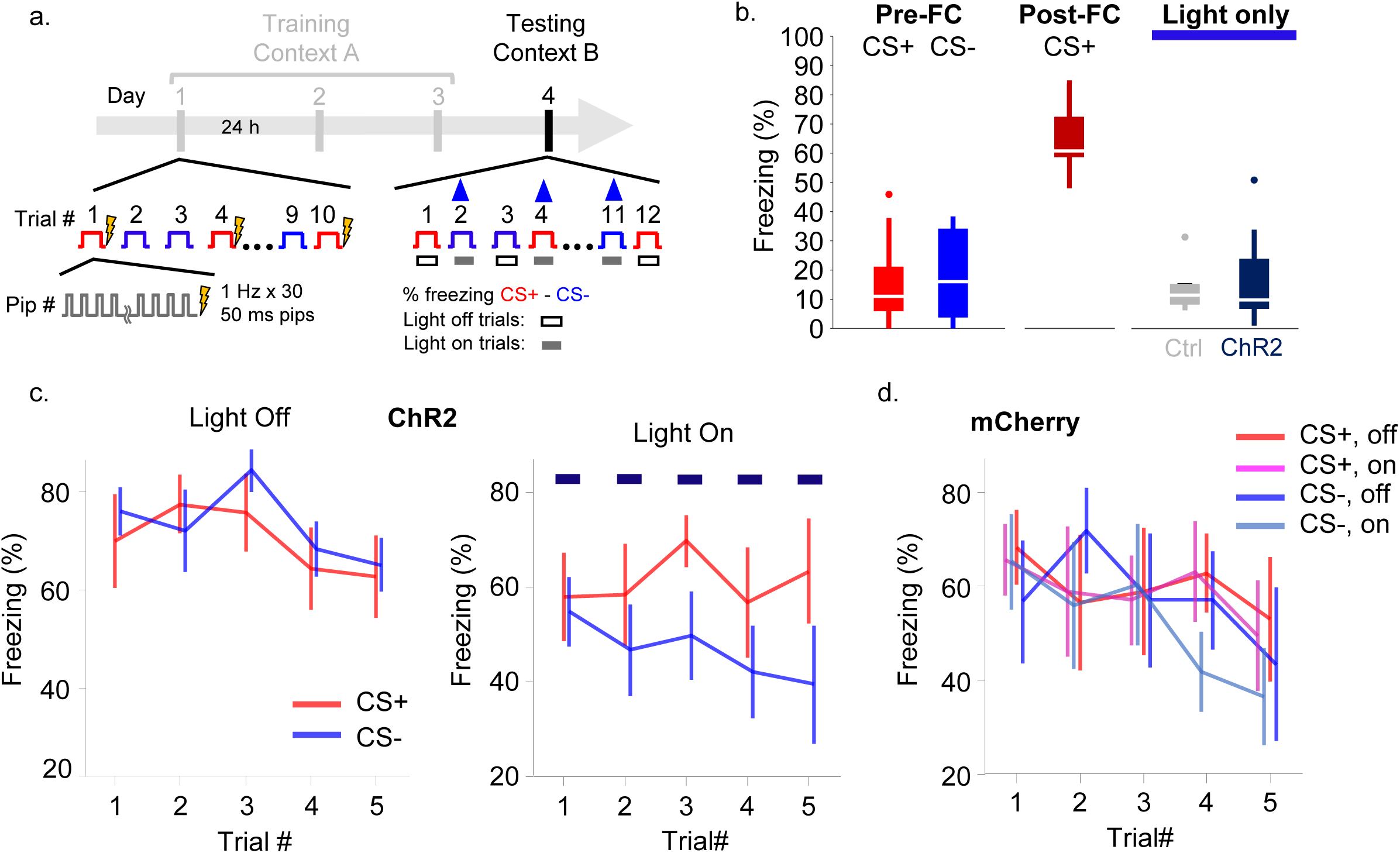
Activation of PL terminals in the BLA during DFC retrieval improves discrimination of the CS-. **a.** The experimental protocol. After 3 days of DFC training, animals undergo DFC retrieval in a new context, when half the CS+ and CS- trials are performed with the Light Off, and half the trials are performed with the Light On to stimulate PL inputs the BLA. **b.** Left, all animals (n=14) showed equivalent low levels of freezing to the CS+ and CS- tones prior to DFC, whereas animals freeze 60-70% of the time to the CS+ (light off) after DFC (n=14, separate data for both groups shown in **Figure 3i-j**). Right, exposure to the light only (in the absence of the CS) had no effect on behavior in either group (Grey, mCherry, n=6; navy, ChR2, n=8). **c**. In the ChR2 group (n=8), animals are freezing equally to the CS+ and CS- on Light Off trials (left panel), whereas they freeze less to the CS- than the CS+ during Light On trials (right panel). **d**. Stimulation of PL terminals in the control group (mCherry, n=6) has no effect on CS+ or CS- retrieval.

## References

1. El-Bar, N., et al., Over-generalization in youth with anxiety disorders. Soc Neurosci, 2017. 12(1): p. 76–85.

2. Hayes, J.P., S.M. Hayes, and A.M. Mikedis, Quantitative meta-analysis of neural activity in posttraumatic stress disorder. Biol Mood Anxiety Disord, 2012. 2: p. 9.

3. Ghosh, S. and S. Chattarji, Neuronal encoding of the switch from specific to generalized fear. Nat Neurosci, 2015. 18(1): p. 112–20.

4. Milad, M.R., et al., Neurobiological basis of failure to recall extinction memory in posttraumatic stress disorder. Biol Psychiatry, 2009. 66(12): p. 1075–82.

5. Grosso, A., et al., A neuronal basis for fear discrimination in the lateral amygdala. Nat Commun, 2018. 9(1): p. 1214.

6. Sangha, S., J.Z. Chadick, and P.H. Janak, Safety encoding in the basal amygdala. J Neurosci, 2013. 33(9): p. 3744–51.

7. Grewe, B.F., et al., Neural ensemble dynamics underlying a long-term associative memory. Nature, 2017. 543(7647): p. 670–675.

8. Shaban, H., et al., Generalization of amygdala LTP and conditioned fear in the absence of presynaptic inhibition. Nat Neurosci, 2006. 9(8): p. 1028–35.

9. Bergado-Acosta, J.R., et al., Critical role of the 65-kDa isoform of glutamic acid decarboxylase in consolidation and generalization of Pavlovian fear memory. Learn Mem, 2008. 15(3): p. 163–71.

10. Wolff, S.B., et al., Amygdala interneuron subtypes control fear learning through disinhibition. Nature, 2014. 509(7501): p. 453–8.

11. Likhtik, E., et al., Prefrontal entrainment of amygdala activity signals safety in learned fear and innate anxiety. Nat Neurosci, 2014. 17(1): p. 106–13.

12. Stujenske, J.M., et al., Fear and safety engage competing patterns of theta-gamma coupling in the basolateral amygdala. Neuron, 2014. 83(4): p. 919–33.

13. Muller, J.F., F. Mascagni, and A.J. McDonald, Postsynaptic targets of somatostatin-containing interneurons in the rat basolateral amygdala. J Comp Neurol, 2007. 500(3): p. 513–29.

14. Meyer, H.C. and D.J. Bucci, The contribution of medial prefrontal cortical regions to conditioned inhibition. Behav Neurosci, 2014. 128(6): p. 644–53.

15. Burgos-Robles, A., et al., Amygdala inputs to prefrontal cortex guide behavior amid conflicting cues of reward and punishment. Nat Neurosci, 2017. 20(6): p. 824–835.

16. Klavir, O., R. Genud-Gabai, and R. Paz, Functional connectivity between amygdala and cingulate cortex for adaptive aversive learning. Neuron, 2013. 80(5): p. 1290–300.

17. Taub, A.H., et al., Oscillations Synchronize Amygdala-to-Prefrontal Primate Circuits during Aversive Learning. Neuron, 2018. 97(2): p. 291–298 e3.

18. Marek, R., et al., Excitatory connections between the prelimbic and infralimbic medial prefrontal cortex show a role for the prelimbic cortex in fear extinction. Nat Neurosci, 2018. 21(5): p. 654–658.

19. Likhtik, E., et al., Prefrontal control of the amygdala. J Neurosci, 2005. 25(32): p. 7429–37.

20. Rosenkranz, J.A. and A.A. Grace, Dopamine attenuates prefrontal cortical suppression of sensory inputs to the basolateral amygdala of rats. J Neurosci, 2001. 21(11): p. 4090–103.

21. Ito, W., B. Fusco, and A. Morozov, Disinhibition-assisted long-term potentiation in the prefrontal-amygdala pathway via suppression of somatostatin-expressing interneurons. Neurophotonics, 2020. 7(1): p. 015007.

